# Double Lock of a Potent Human Monoclonal Antibody against SARS-CoV-2

**DOI:** 10.1101/2020.11.24.393629

**Authors:** Ling Zhu, Yong-Qiang Deng, Rong-Rong Zhang, Zhen Cui, Chun-Yun Sun, Chang-Fa Fan, Xiaorui Xing, Weijin Huang, Qi Chen, Na-Na Zhang, Qing Ye, Tian-Shu Cao, Nan Wang, Lei Wang, Lei Cao, Huiyu Wang, Desheng Kong, Juan Ma, Chunxia Luo, Yanjing Zhang, Jianhui Nie, Yao Sun, Zhe Lv, Neil Shaw, Qianqian Li, Xiao-Feng Li, Junjie Hu, Liangzhi Xie, Zihe Rao, Youchun Wang, Xiangxi Wang, Cheng-Feng Qin

## Abstract

Receptor recognition and subsequent membrane fusion are essential for the establishment of successful infection by SARS-CoV-2. Halting these steps can cure COVID-19. Here we have identified and characterized a potent human monoclonal antibody, HB27, that blocks SARS-CoV-2 attachment to its cellular receptor at sub-nM concentrations. Remarkably, HB27 can also prevent SARS-CoV-2 membrane fusion. Consequently, a single dose of HB27 conferred effective protection against SARS-CoV-2 in two established mouse models. Rhesus macaques showed no obvious adverse events when administrated with 10-fold of effective dose of HB27. Cryo-EM studies on complex of SARS-CoV-2 trimeric S with HB27 Fab reveal that three Fab fragments work synergistically to occlude SARS-CoV-2 from binding to ACE2 receptor. Binding of the antibody also restrains any further conformational changes of the RBD, possibly interfering with progression from the prefusion to the postfusion stage. These results suggest that HB27 is a promising candidate for immuno-therapies against COVID-19.

**Highlights:**

1. SARS-CoV-2 specific antibody, HB27, blocks viral receptor binding and membrane fusion
2. HB27 confers prophylactic and therapeutic protection against SARS-CoV-2 in mice models
3. Rhesus macaques showed no adverse side effects when administered with HB27
4. Cryo-EM studies suggest that HB27 sterically occludes SARS-CoV-2 from its receptor

## Introduction

On March 11^th^ 2020, the World Health Organization declared the 2019 coronavirus disease (COVID-19) as a pandemic. Severe acute respiratory syndrome coronavirus 2 (SARS-CoV-2), the etiological agent of this pandemic continues to ravage the global population, causing millions of infections. Losses in lives, declining wellbeing, and disruption of economic activities as a result of the infections have strained societies and significant impacted on people’s normal life. SARS-CoV-2 belongs to the betacoronavirus genus, five coronaviruses of which, together with two alphacoronaviruses, endowed with an ability to infect humans (Lu et al., 2020; Zhou et al., 2020). Among these, infections caused by SARS-CoV, SARS-CoV-2 and Middle East Respiratory Syndrome coronavirus (MERS-CoV) are known to culminate into more severe clinical manifestations (Gao et al., 2020). To date, no specific drugs or vaccines effective against these highly pathogenic coronaviruses have been approved.

Like, SARS-CoV, SARS-CoV-2 utilizes its protuberant S glycoprotein to engage with its cellular receptor, human angiotensin converting enzyme 2 (ACE2), for forging membrane fusion in order to enter host cell (Gallagher and Buchmeier, 2001; Hoffmann et al., 2020). Each monomeric S protein can be cleaved by host proteases, such as TMPRSS2 (Hoffmann et al., 2020; Shang et al., 2020) into two functional domains, the distal globular S1 domain and the membrane-proximal S2 domain, which mediate receptor binding and membrane fusion, respectively (Li, 2016). The S1 subunit consists of an N-terminal domain (NTD) and a C-terminal domain, which often functions as the receptor binding domain (RBD). Conformational transitions are triggered upon release of the S1 subunit after receptor binding and subsequent priming of the protein by host cell proteases. These two key events advance the life-cycle of the virus from the prefusion to the postfusion stage, leading to the fusion of the viral membrane with that of the host cell (Li, 2016; Walls et al., 2017).

Such important roles played by S during viral infection make them valuable targets for antibody-based drug and vaccine design (Pallesen et al., 2017). Previous structural studies have revealed that the S trimer can switch between a receptor-accessible state where one or more RBDs are in the open conformation and a receptor-inaccessible state where all the RBDs are in the closed conformation. This switch is accomplished through a hinge-like movement of the RBD, indicative of a dynamic and complicated protein-protein interaction mode with host cells (Gui et al., 2017; Kirchdoerfer et al., 2016; Walls et al., 2020; Wrapp et al., 2020; Zhe Lv, 2020). Although numerous neutralizing antibodies (NAbs) targeting the RBDs of SARS-CoV or MERS have been reported (Corti et al., 2015; Du et al., 2009; Walls et al., 2019), the immunogenic features and key epitopes of SARS-CoV-2 remain poorly characterized. Recently, a cross-binding mAb, CR3022, was demonstrated to neutralize SARS-CoV, but it failed to efficiently prevent SARS-CoV-2 infection, highlighting the challenges posed by conformationally flexible virus-specific neutralizing epitopes in conferring protection against infection (Yuan et al., 2020). More recently, a number of NAbs have been shown to block the binding of SARS-CoV-2 to ACE2 and another RBD-targeting NAb, S309, acted by inducing antibody-dependent cell cytotoxicity (ADCC) which surprisingly did not involve the blocking of virus-receptor interaction (Pinto et al., 2020; Wu et al., 2020). This raises the possibility of existence of hitherto undiscovered neutralization mechanisms for SARS-CoV-2 RBD-targeting NAbs. A detailed understanding of the mechanisms underlying the neutralization of SARS-CoV-2 is likely to help provide new guidance for the development of antiviral therapeutics and rational vaccine design.

## Results

### Phage display identifies a potent SARS-CoV-2 specific NAb

We previously identified a set of NAbs from an antibody library which was generated from RNAs extracted from peripheral lymphocytes of mice immunized with recombinant SARS-CoV RBD protein (Zhe Lv, 2020). In this study, we constructed another antibody library by immunizing mice with recombinant SARS-CoV-2 RBD, which yielded a chimeric anti-SARS-CoV-2 mAb, named mhB27. mhB27 was able to strongly bind to SARS-CoV-2 RBD and exhibited potent neutralizing activities against SARS-CoV-2 when tested in a vesicular stomatitis virus (VSV) pseudotyping system (PSV) (Figure S1). A humanized antibody HB27 was generated based on the sequences of mhB27. To investigate the viral specificity of HB27, we performed binding assays measuring the ability of HB27 to bind the RBDs of SARS-CoV, SARS-CoV-2 and MERS-CoV. Analysis of the data obtained from real-time quantitation and kinetic characterization of biomolecular interactions using OCTET system demonstrated that both immunoglobulin G (Ig G) and Fab fragments of HB27 bind tightly to SARS-CoV-2 RBD with affinities of 0.07 nM and 0.27 nM, respectively. However, this antibody exhibits undetectable interactions with the RBDs of SARS-CoV and MERS-CoV, suggesting that HB27 is SARS-CoV-2-specific (Figure 1A-1C). HB27 showed potent neutralizing activities against SARS-CoV-2 with a 50% inhibition concentration (IC_50_) value of 0.04 nM. Perhaps correlated with the inability to interact with SARS-CoV RBD, HB27 possessed no inhibition activity against SARS-CoV in PSV-based neutralization assays (Figure 1D-1E). Classical plaque reduction neutralization test (PRNT) conducted against an authentic SARS-CoV-2 strain (BetaCoV/Beijing/IME-BJ01/2020) further verified its neutralizing activity with a PRNT_50_ value of 0.22 nM (Figure 1F).

**Figure 1.**
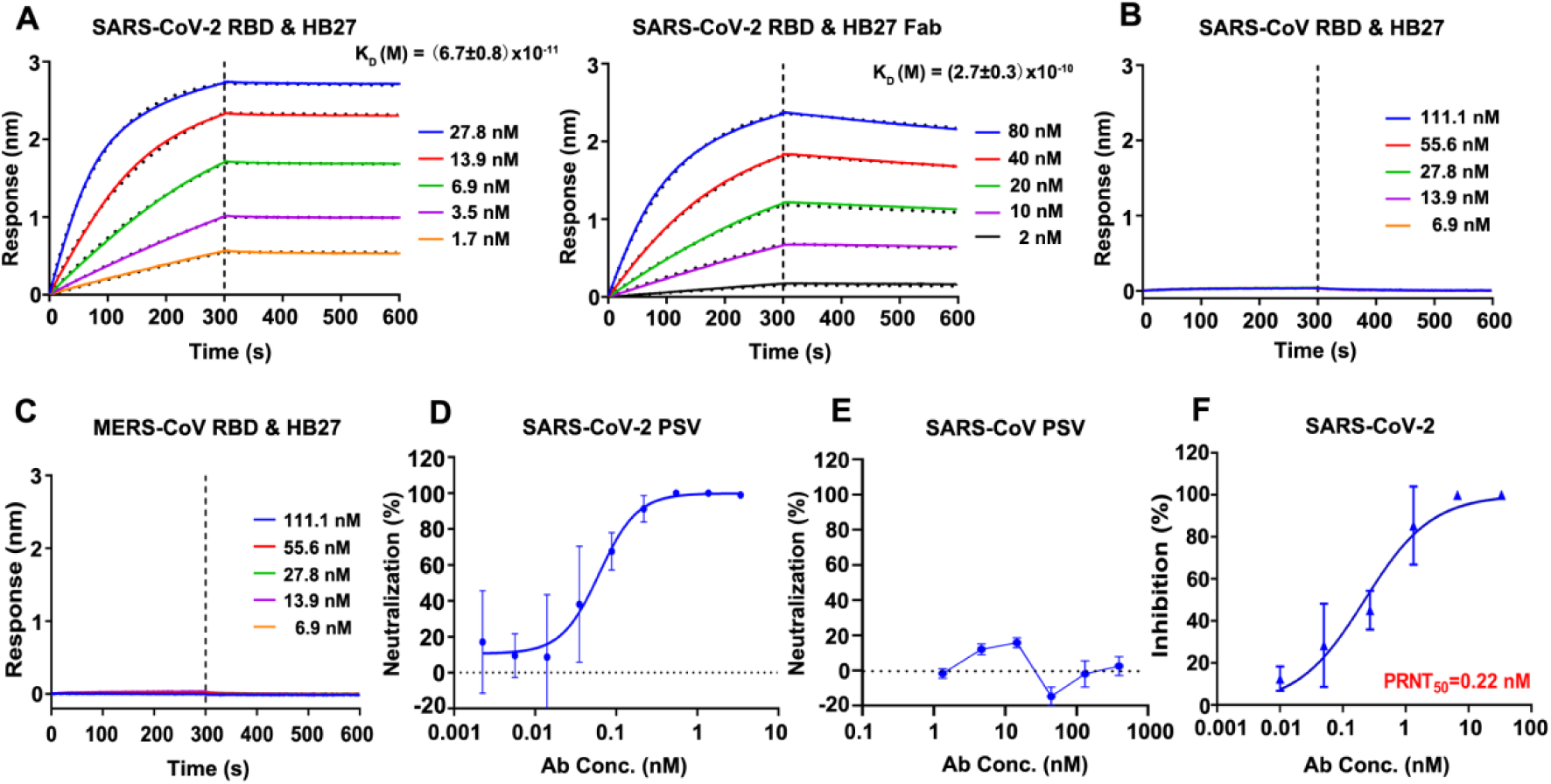
HB27 is a SARS-CoV-2-specific antibody of high potency. (A) Analysis of affinity of HB27 (left panel) and HB27 Fab fragments (right panel) for SARS-CoV-2 RBD. Biotinylated SARS-CoV-2 RBD protein was loaded on Octet SA sensor and tested for real-time association and dissociation from HB27 IgG and HB27 Fab fragments, respectively. (B) and (C) Analysis of affinity of HB27 for SARS-CoV RBD and MERS-CoV RBD, respectively. (D) and (E) Neutralizing activity of HB27 against SARS-CoV-2 and SARS-CoV pseudoviruses (PSV), respectively. Serially diluted HB27 titres were added to test neutralizing activity against SARS-CoV-2 and SARS-CoV PSV. (F) *In vitro* neutralization activity of HB27 against SARS-CoV-2 by plaque reduction neutralization test (PRNT) in Vero cells. Neutralizing activities are represented as mean ± SD. Experiments were performed in duplicates See also Figure S1.

### Prophylactic and therapeutic efficacy of HB27 in SARS-CoV-2 susceptible mice

Given the excellent neutralizing activities at sub-nM concentrations, we next sought to assess the correlation between *in vitro* neutralization and *in vivo* protection. The HB27 produced in the CHO cell line was first tested in a newly established mouse model based on a SARS-CoV-2 mouse adapted strain MASCp6 (Gu et al., 2020). Upon MASCp6 intranasal challenge, adult BALB/c sustained robust viral replication in the lungs at 3-5 days post inoculation. To evaluate the protection efficacy of HB27, BALB/c mice challenged with MAScp6 were administered a single dose of 20 mg/kg of HB27 in prophylactic as well as therapeutic settings (Figure 2A). As expected, high levels of viral RNAs were detected in the lungs and trachea at 3 and 5 days post infection in the control group of mice treated with PBS (Figure 2B-2C). Remarkably, a single dose of HB27 administered either before or post SARS-CoV-2 exposure resulted in >99.9% reduction of the viral RNA loads in the lungs and trachea (Figure 2B-2C).

**Figure 2.**
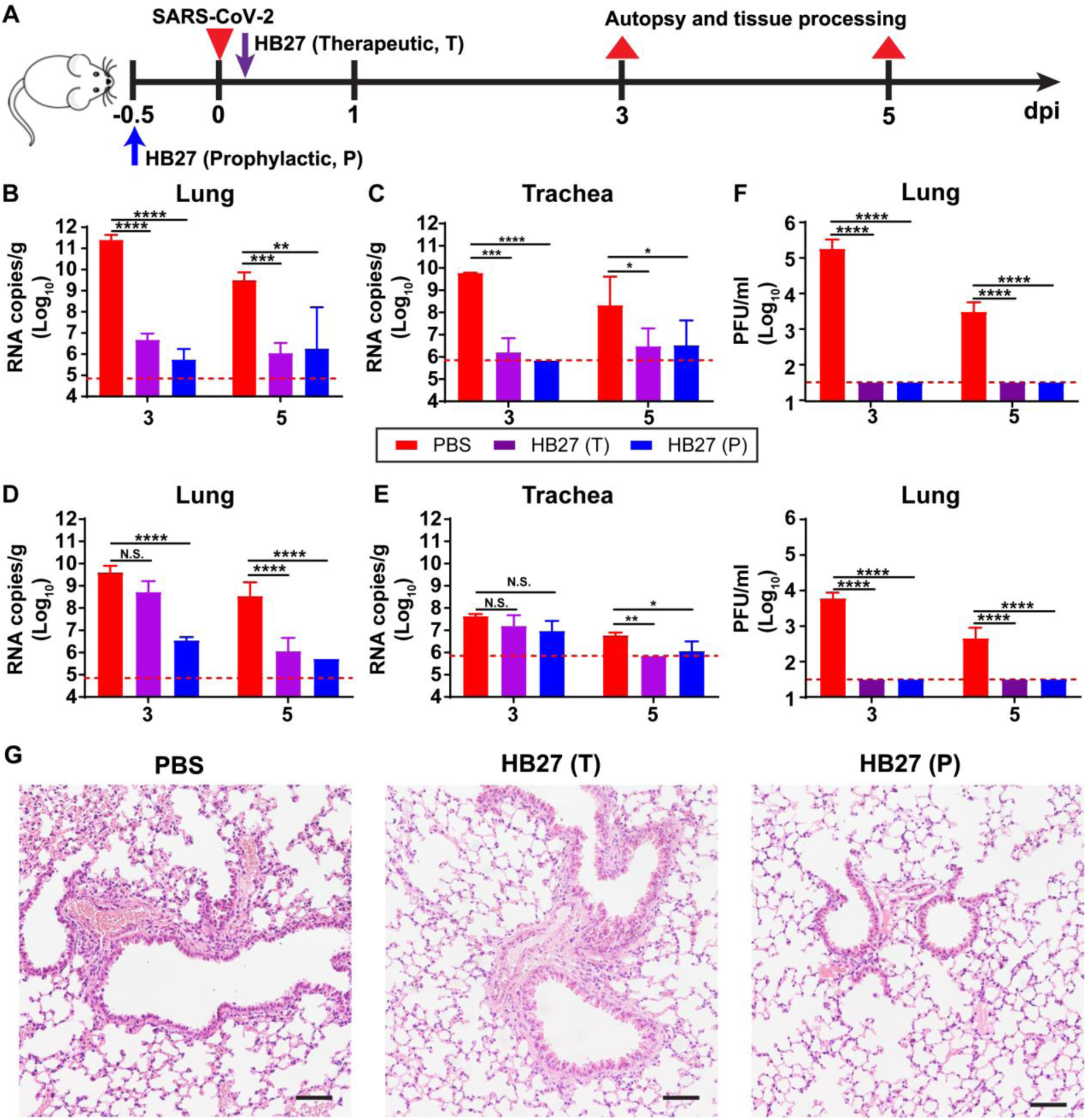
Prophylactic and therapeutic efficacy of HB27 in two SARS-CoV-2 susceptible mice models. (A) Experimental design for therapeutic and prophylactic evaluations of HB27 in two SARS-CoV-2 susceptible mice models. Group of 6-to-8 week-old hACE2 mice and BALB/c mice were infected intranasally with 5×10^4^ PFU of SARS-CoV-2 BetaCoV/Beijing/IME-BJ01/2020 or 1.6×10^4^ PFU of MASCp6 as described previously, respectively. A dose of 20 mg/kg HB27 was injected intraperitoneally at 12 hours before infection (the prophylactic group, P) or at 2 hours after infection (the therapeutic group, T). PBS injections were used as control group. Then, the lung tissues of mice were collected at 3 and 5 dpi for virus titer, H&E and Immunostaining. (B) and (C) Virus titers of lung and trachea tissues at 3 or 5 dpi in mouse model based on a SARS-CoV-2 mouse adapted strain MASCp6. The viral loads of the tissues were determined by qRT-PCR (*P< 0.05; **P< 0.01; ***P< 0.001; ****P< 0.0001; n.s., not significant). Data are represented as mean ± SD. Dashed lines represents limit of detection. (D) and (E) Virus titers of lung and trachea tissues at 3 or 5 dpi in hACE2 humanized mouse model. The viral loads of the tissues were determined by qRT-PCR (*P< 0.05; **P< 0.01; ***P< 0.001; ****P< 0.0001; n.s., not significant). Data are represented as mean ± SD. Dashed lines represents limit of detection. (F) Viral burden at 3 or 5 dpi in the lungs from two mouse models (up: BALB/c mice; bottom: hACE2 mice), measured by plaque assay. Data are represented as mean ± SD. Dashed lines represents limit of detection. (G) Histopathological analysis of lung samples at 5 dpi. Scale bar: 100 µm.

Furthermore, we validated the *in vivo* protection efficacy of HB27 in a human ACE2 (hACE2) humanized mouse model that was susceptible to SARS-CoV-2 infection (Sun et al., 2020). Similar to the studies with the MASCp6 strain of mice, either prophylactic or therapeutic administration of HB27 conferred a clear benefit on the hACE2 humanized mouse model as indicated by a significant reduction in viral RNA loads in the lungs and trachea at day 5 post SARS-CoV-2 challenge. Prophylactic administration of HB27 showed a more potent antiviral effect, resulting in >1,000-fold reduction in lung viral levels (Figure 2D-2E). A direct challenge via administration of excessive (up to 5 × 10^5^ PFU) SARS-CoV-2 through the intranasal route, where the IgG antibodies may not be able to directly engage the target, could lead to virus particles gaining access to the lung and trachea. Such experimental observations in the prophylactic and therapeutic settings for many other protective human antibodies against SARS-CoV-2 have been reported (Zhe Lv, 2020; Zost et al., 2020). However, it’s worthy to note that no infectious virions could be detected in the lung at day 3 and day 5 as measured by a plaque assay of lung tissue homogenates (Figure 2F). These results suggest that the low levels of viral RNA copies detected in the lung/trachea might be the remnants of the viral genomes from the infection at the very early stage. Histopathological examination revealed moderate interstitial pneumonia, characterized by inflammatory cell infiltration, alveolar septal thickening and distinctive vascular system injury developed in hACE2 humanized mice belonging to the PBS control group at day 5 (Figure 2G). In contrast, the lungs in mice from the HB27 treated group only showed minimal or very mild inflammatory cell infiltration, and no obvious lesions of alveolar epithelial cells or focal hemorrhage (Figure 2G). Collectively, these results clearly demonstrated the utility of HB27 for prophylactic or therapeutic purposes against SARS-CoV-2.

### Evaluation of the safety of administration of HB27 in non-human primates

As part of the non-clinical safety studies prior to the conduction of human clinical trials for pharmaceuticals, we systematically evaluated the safety of administration of HB27 in rhesus macaques. Two groups of 4 animals (n=4) were administered a single intravenous high dose (500 mg/kg, 10-fold of estimated effective dose in human) of HB27 or placebo. HB27 serum concentrations, clinical observations and biological indices were monitored closely for 16 days (Figure 3). Neither fever nor weight loss was observed in any macaque after the infusion of HB27, and the appetite and mental state of all animals remained normal. The toxicokinetics of HB27 in rhesus macaques was evaluated by measuring HB27 levels in serum pre-infusion and at indicated time intervals after administration. A mean maximum serum concentration (Cmax) of 12.8 mg/mL (± 0.8) of HB27 could be achieved and the average half-life of the antibody was 10.0 days (± 2.2) (Figure 3A and Table S1). Notably, prophylactic and therapeutic efficacy of HB27 in animal models revealed that >99.9% of the viral RNA loads in the lungs and trachea could be reduced at 5 days post infection (Figure 2), suggesting that a half-life of 10 days for HB27 is probably sufficient for deriving therapeutic benefit. Details of the results of the measurements of toxicokinetic parameters are presented in Table S1. These results suggest that HB27 probably has pharmacokinetic properties consistent with a typical human IgG1. Hematological and biochemical analysis, including biochemical blood tests and lymphocyte subset percent (CD4^+^ and CD8^+^) showed no notable changes in the HB27 administrated group when compared to the placebo group (Figure 3B-3D). Taken together, the results of our animal studies indicate that HB27 is generally safe in non-human primates.

**Figure 3.**
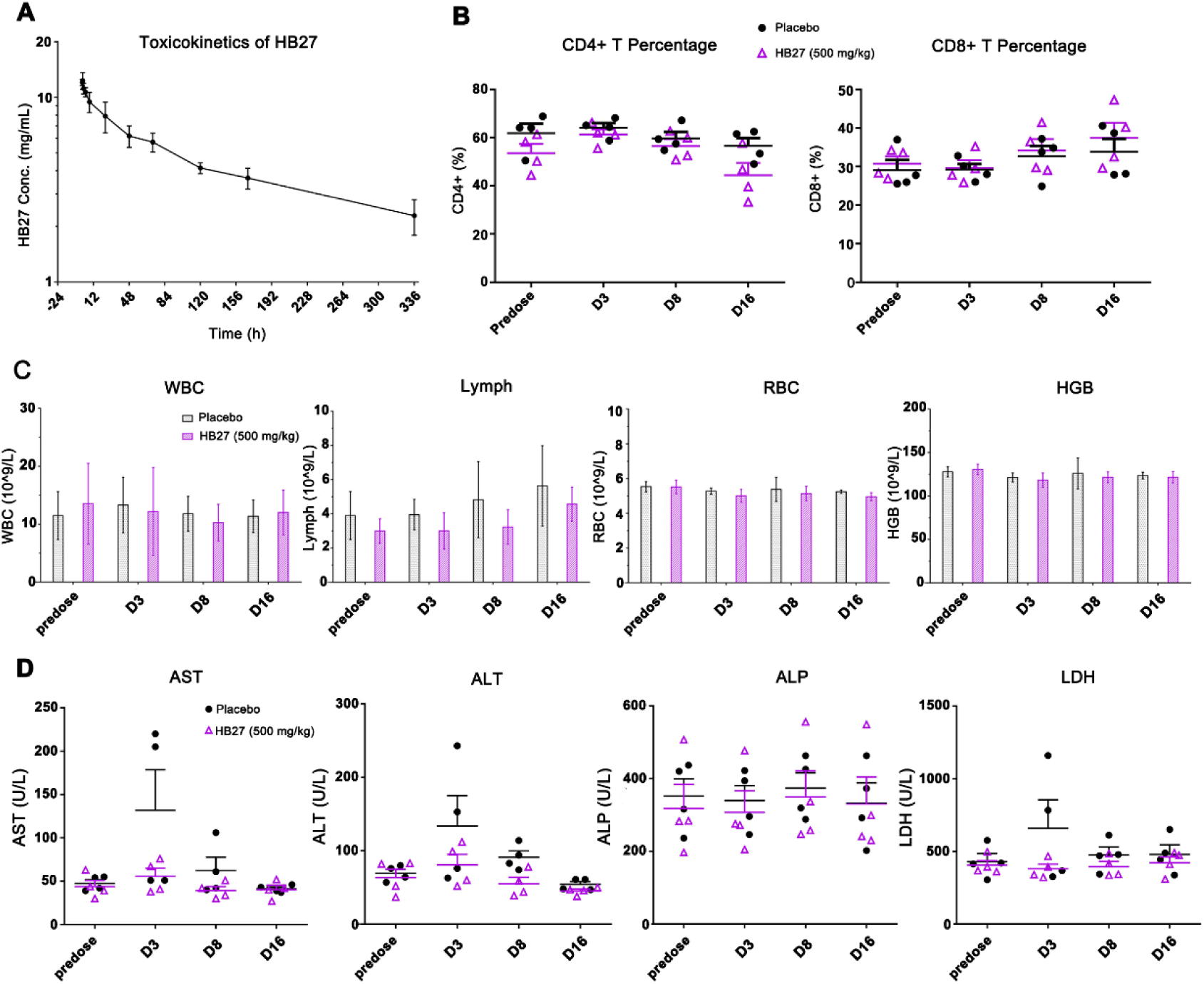
Safety evalution of HB27 in rhesus macaques. (A) The toxicokinetics of HB27 in rhesus macaques was evaluated by measuring HB27 levels in serum predose and at 12, 48, 84, 120, 156h, 192, 228, 264, 300 and 336 hours after administration. (B-D) Rhesus macaques were given intravenous injections of a single dose of either placebo or HB27 (500 mg/kg), and monitored by lymphocyte subset analysis (B), hematological test (C), and biochemical blood test (D) predose and 3, 8 and 16 days postdose. WBC: white blood cells; Lymph: lymphocytes; RBC: red blood cells; HBG: hemoglobin; AST: aspartate transaminase; ALT: alanine transaminase; ALP: alkaline phosphatase; LDH: lactate dehydrogenase See also Table S1.

### HB27 prevents the attachment of SARS-CoV-2 to host cells by blocking its binding to ACE2

To evaluate the ability of HB27 to inhibit binding of RBD to ACE2, we performed competitive binding assays at both protein and cellular levels. Results of the enzyme-linked immunosorbent assay (ELISA) revealed that HB27 could prevent the binding of soluble ACE2 (monomer in solution) to SARS-CoV-2 RBD with an EC_50_ value of 0.5 nM (Figure S2A). To verify the ability of HB27 to block the binding of ACE2 to trimeric S, we expressed and purified stabilized SARS-CoV-2 S ectodomain trimer. Surface plasmon resonance (SPR) assays indicated that HB27 interacts with SARS-CoV-2 S trimer with a slightly stronger binding affinity (∼0.04 nM) (Figure S2B), which was about 1000-fold higher than that of soluble ACE2 with SARS-CoV-2 S (Figure S2C) (Shang et al., 2020). For the competitive SPR, two sets of assays: exposing the trimeric S to HB27 first and then to soluble ACE2, or the other way around, were conducted. As expected, binding of HB27 completely blocked the interaction between soluble ACE2 and SARS-CoV-2 trimeric S. Moreover, soluble ACE2 that had already bound to trimeric S could be replaced by HB27 because of the ∼1000-fold difference in binding affinities of these ligands to the SARS-CoV-2 trimeric S (Figure 4A). Cell-based immunofluorescent blocking assays demonstrated that HB27 could block both the binding of soluble ACE2 to SARS-CoV-2 S expressing 293T cells and the attachment of SARS-CoV-2 RBD to ACE2 expressing 293T cells in a dose dependent manner albeit with relative high EC_50_ values of about 5-50 nM (Figure 4B and Figure S2D). Overexpression of ACE2/ S trimer on the 293T cell surface and the presence of the dimeric form of ACE2 on cell surface are probably the reasons for the substantially higher concentration of HB27 needed to prevent attachment of the virus to the cell surface. To further verify these results in cell-based viral infection model, we used real-time reverse transcriptase–PCR (RT–PCR) to quantify the amount of virus remaining on the surface of cells that were treated with HB27 pre- and post-viral attachment at 4 °C. In line with the results of the competitive binding assays, HB27 efficiently prevented SARS-CoV-2 attachment to host cell surface at sub-nM and could displace the virions that had already bound to the cell surface at ∼2.5 nM (Figure 4C).

**Figure 4.**
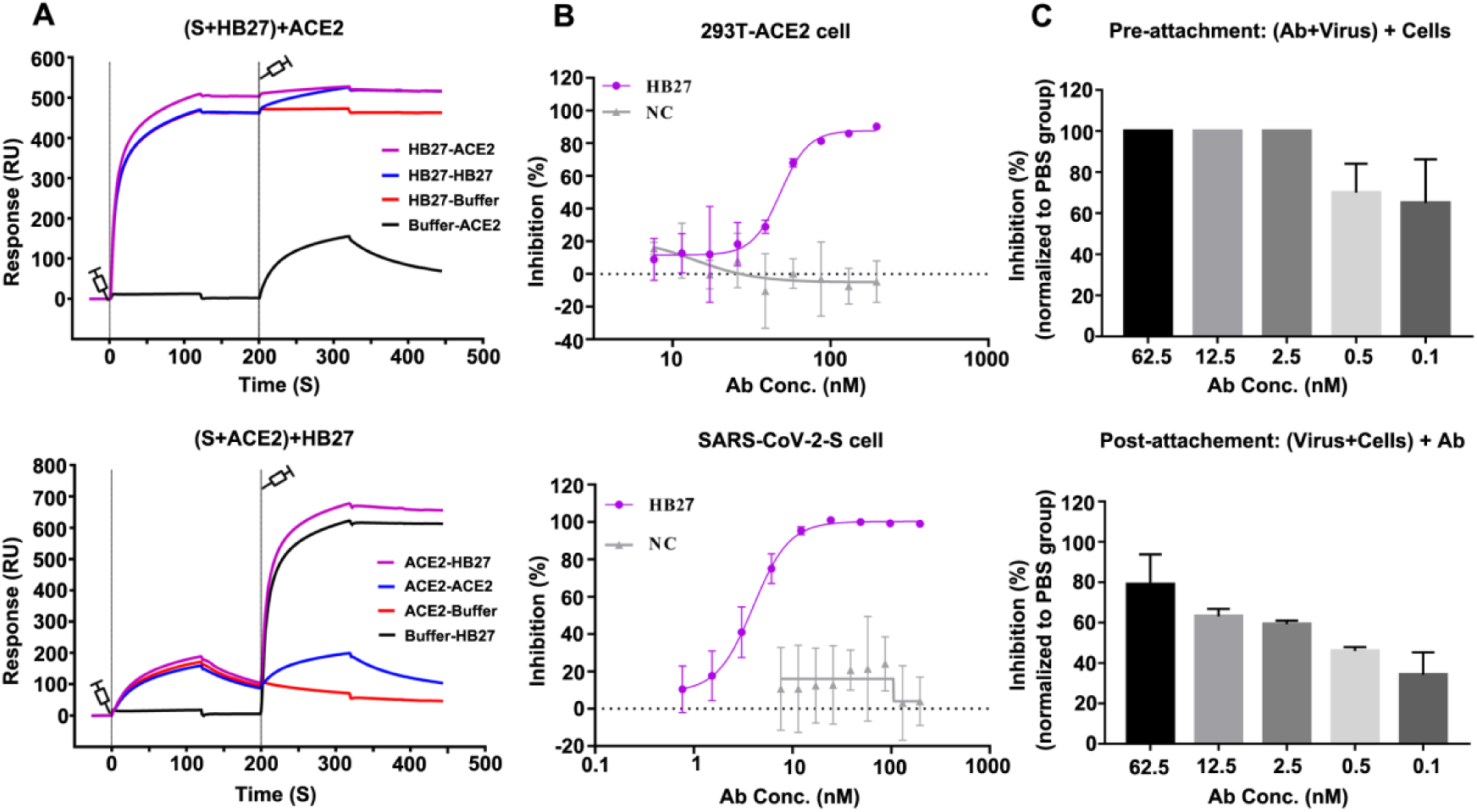
HB27 blocks the interactions of SARS-CoV-2 with ACE2. (A) BIAcore SPR kinetics showing the competitive binding of HB27 and ACE2 to SARS-CoV-2 S trimer. For both panels, SARS-CoV-2 S protein was immobilized onto the sensor chips. In the upper panel, HB27 was first injected, followed by ACE2, whereas in the lower panel, ACE2 was injected first and then HB27. The control groups are as shown by the curves. (B) Blocking of SARS-CoV-2 RBD binding to 293T-ACE2 cells by HB27 (upper panel). Recombinant SARS-CoV-2 RBD protein and serially diluted HB27 were incubated with ACE2 expressing 293T cells (293T-ACE2) and tested for binding of HB27 to 293T-ACE2 cells. Competitive binding of HB27 and ACE2 to SARS-CoV-2-S cells (lower panel). Recombinant ACE2 and serially diluted HB27 were incubated with 293T cells expressing SARS-CoV-2 S (SARS-CoV-2-S) and tested for binding of HB27 to SARS-CoV-2-S cells. BSA was used as a negative control (NC). (C) Amount of virus on the cell surface, as detected by RT-PCR, when exposed to HB27 prior to (upper panel) and after (lower panel) the virus was allowed to attach to cells. Values are mean ± SD. Experiments were repeated in triplicate. See also Figure S2.

### HB27 prevents SARS-CoV-2 membrane fusion

A common way to determine whether the antibody inhibits virus-receptor binding or a post-attachment step of the infection is to compare neutralization curves deduced from mixing antibody with the virus before or after binding to cells at 4 °C. The assumption is antibodies that inhibit receptor binding will not have a neutralizing effect on virus that has already bound to its receptor. However, this assumption may not be true, because high-affinity antibodies could possibly displace the virus that is already complexed to a low-affinity receptor, as observed for HB27 (Fig. 4B-4C). Thus, deriving the mechanism of neutralization by just conducting neutralization assays may not yield the complete picture, although pre- and post-attachment neutralization assays suggested that HB27 inhibits a post-attachment step of the infection (Figure 5A). In coronaviruses, receptor binding and proteolytic processing act in synergy to trigger a series of conformational changes in S, bringing viral and cellular membranes in proximity for fusion, leading to establishment of infection (Walls et al., 2017). TMPRSS2-mediated cleavage is capable of activating the fusion potential of coronavirus S proteins, inducing receptor-dependent syncytium formation, which was recently observed in natural SARS-CoV-2 infections as well (Ou et al., 2020; Xia et al., 2020). To explore whether HB27 could interfere with syncytium formation, we established the S-mediated cell-cell fusion system using 293T cells that express SARS-CoV-2 S with a GFP tag as the effector cells and Vero-E6 cells as the target cells (Figure 5B). After co-incubation of effector and target cells for 48 h, hundreds of cells fused together into one large syncytium with multiple nuclei (Figure 5B). Remarkably, HB27 could completely inhibit SARS-CoV-2 mediated cell-cell fusion at the concentration of 0.5 μM. Notably, this result is comparable with the inhibition efficiencies of some pan-coronavirus fusion inhibitors (Figure 5B) (Xia et al., 2020). Neither SARS-CoV-2 RBD-targeting neutralizing antibody, H014 nor the isotype control antibody (anti-H7N9) could prevent membrane fusion under similar conditions (Figure 5B). Furthermore, we performed live SARS-CoV-2 neutralization assay in a post-binding manner in Huh7 cells. Briefly, Huh7 cells were infected with 100 PFU of SARS-CoV-2 for 1 h at 4 °C. Unbound viral particles were washed away using buffer. After that cells were further cultured in the presence of a series of concentrations (0, 4, 20 and 100 nM) of HB27, or 100 nM of H014 at 37 °C for 48 h. Similar to S-mediated cell-cell fusion, the large syncytiums formed by live SARS-CoV-2 infected Huh7 cells were observed in the absence of HB27 and the presence of 100 nM H014 (Figure 5C). Expectedly, HB27 could significantly inhibit SARS-CoV-2 mediated formation of the syncytiums in a dose dependent fashion and completely block the cell-cell fusion at 100 nM (Figure 5C). Notably, such inhibition, to some extent, can possibly be attributed to the ability of HB27 to strip SARS-CoV-2 off the cell surface. To further characterize the molecular basis for fusion inhibition by HB27, we established an *in vitro* membrane fusion assay that treatments of purified SARS-CoV-2 virions by trypsin and ACE2 could trigger viral membrane fusion with liposome at acidic environment. Liposome fusion results show that HB27, but not H014, is capable of efficiently blocking pH-dependent fusion of SARS-CoV-2 with liposomes in a dose dependent manner (Figure 5D). The blockage of membrane fusion by HB27 is likely another important mechanism of neutralization. However, given the relatively higher concentration of HB27 needed to block viral membrane fusion, blocking viral attachment to its host cell receptor is likely to be the main mechanism of neutralization.

**Figure 5.**
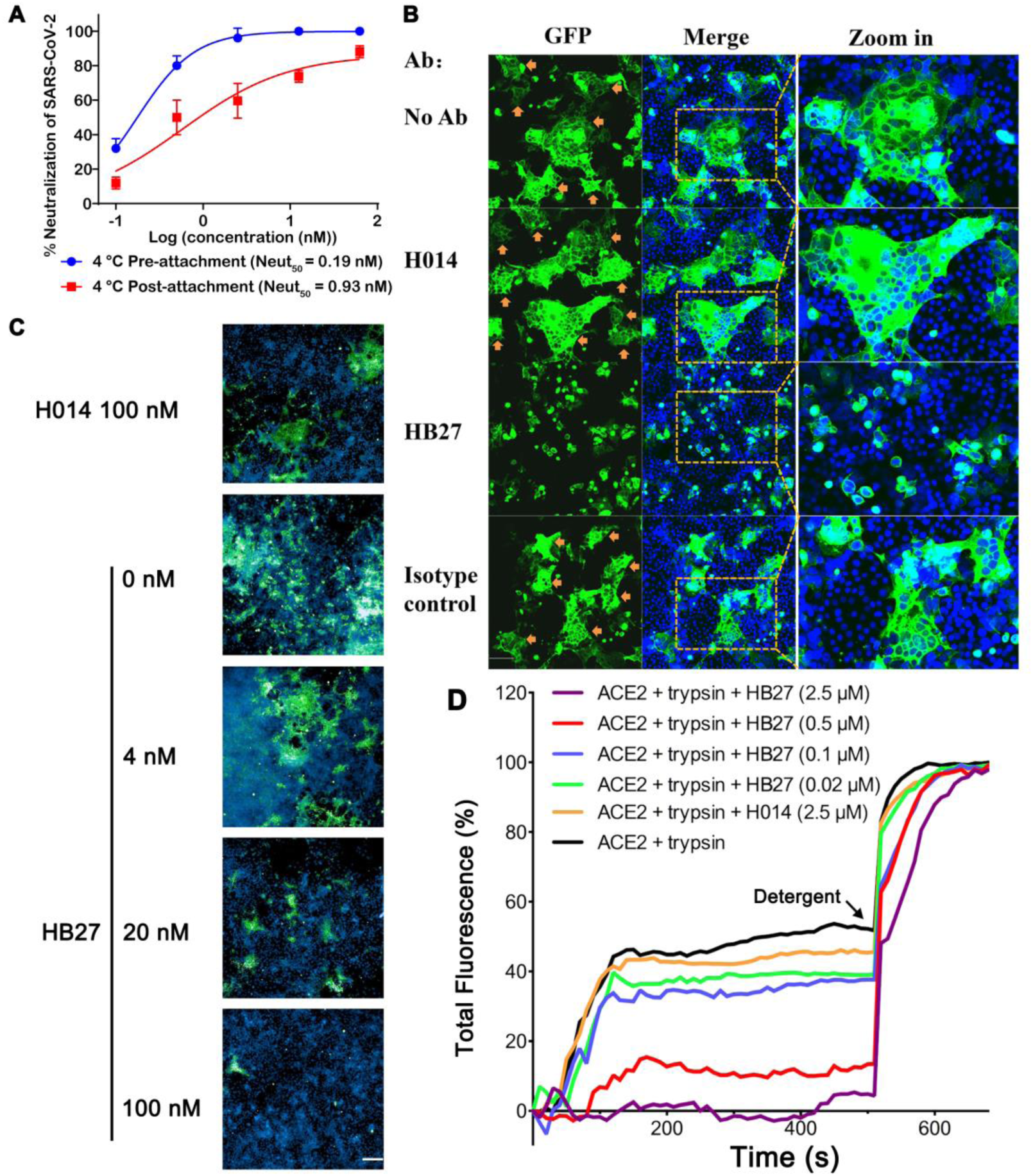
HB27 inhibits SARS-CoV-2 membrane fusion. (A) HB27 had potent neutralization activities when exposed to virus before or after attachment to Huh7 cells. Values are mean ± SD. Experiments were repeated in triplicate. (B) HB27 inhibits S protein-mediated cell-cell fusion. 293T cells were transfected with SARS-CoV-2 S-GFP protein, co-cultured with Vero E6 cells in the absence or presence of 100 μg/mL H014 or HB27 or anti-influenza H7N9 antibody (isotype control). No Ab: in the absence of antibodies. Images were taken after 48 h. Cells were fixed with 4% paraformaldehyde (PFA) at room temperature for 20 min and stained for nuclei with 4,6-diamidino-2-phenylindole (DAPI). (C) HB27 inhibits SARS-CoV-2-mediated cell-cell fusion. Huh7 cells were infected with 100 PFU of SARS-CoV-2 for 1 h at 4°C and washed for 3 times. After that cells were further cultured in the presence of a series of concentrations (0, 4, 20 and 100 nM) of HB27, or 100 nM of H014 at 37 °C for 48 h. Images were taken after 48 h. Cells were fixed with 4% (w/v) PFA for 20 min and incubated with anti-SARS-CoV-2 S protein antibody and stained for nuclei with DAPI. Scale bar equals 200 μm. (D) HB27 blocks receptor-mediated fusion of SARS-CoV-2 with liposomes. Liposomes were loaded with self-quenching concentrations of the fluorescent dye calcein. Perturbation of the bilayer causes the release of calcein resulting in dilution and a consequent increase in its fluorescence. Fusion of SARS-CoV-2 with liposomes occurred in the presence of both ACE2 and trypsin and a series of HB27 concentrations were used to inhibit the fusion. 10% Triton X-100 treatment was used to achieve 100% calcein leakage. All data shown are representative of three independent experiments.

### Structural basis for the SARS-CoV-2 specific binding of HB27

To delineate the structural basis for HB27-mediated specific neutralization, we determined the cryo-EM structure of a prefusion stabilized SARS-CoV-2 S ectodomain trimer in complex with the HB27 Fab fragment using single particle reconstruction. Similar to previously published studies on *apo* SARS-CoV-2 S trimer, two distinct conformational states referred to as the “close” and “open” RBDs were observed in the structure of the complex (Figure 6A). Cryo-EM characterization of the complex showed full occupancy with one Fab bound to each RBD of the homotrimeric S. We asymmetrically reconstructed the complex structure at an overall resolution of 3.5 Å, which represents two “open” and one “close” RBDs (Figure 6A, Figure S3-S4 and Table S2). The initial maps for the binding interface between RBD and HB27 were relatively weak due to conformational heterogeneity, which is in line with the structural observations of stochastic RBD rotations at different angles while switching from the “closed” to “open” states (Figure 6B-6C). In order to improve the local resolution, we employed a “block-based” reconstruction approach, which resulted in a 3.9 Å resolution, enabling reliable analysis of the interaction interface (Figure S3-S4).

**Figure 6.**
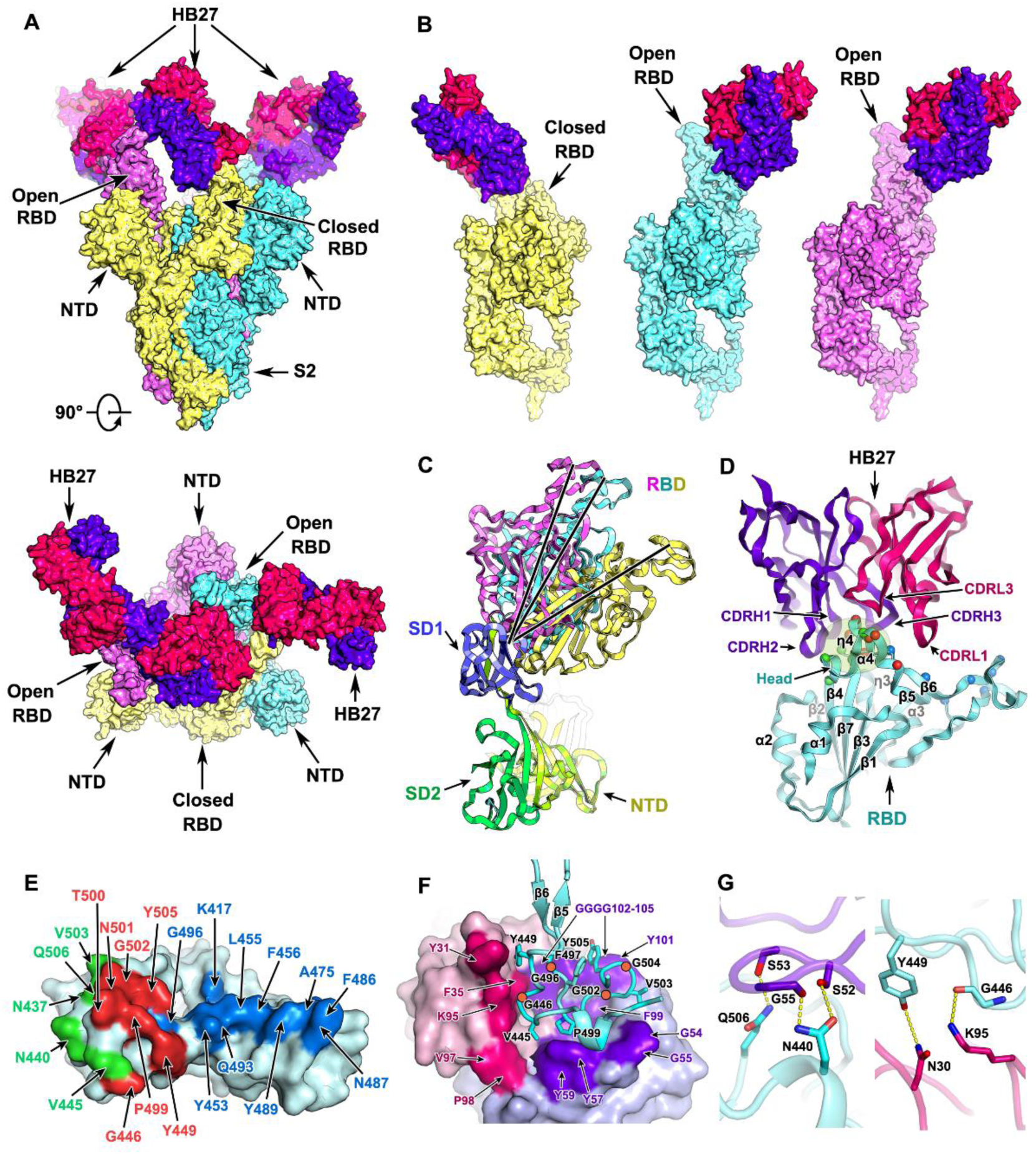
Structure and interaction of the SARS-CoV-2 S trimer with HB27. (A) Orthogonal views of SARS-CoV-2 S trimer in complex with three copies of HB27 Fab. (B) Individual views of the three monomers each complexed with one HB27 Fab. (A) and (B) The S trimer and HB27 are rendered as molecular surfaces. Three monomers of the S trimer are colored in yellow, cyan and violet, respectively. The HB27 light and heavy chains are colored in hotpink and purpleblue, respectively. RBD: receptor binding domain. NTD: N-terminal domain. S2: the S2 subunit. (C) S1 subunits of the three monomers from SARS-CoV-2 S trimer complexed with HB27 are superposed; HB27 Fabs are not shown. All domains are presented as ribbon diagrams. Three RBD domains are colored in yellow, cyan and violet, respectively. SD1: subdomain 1. SD2: subdomain 2. (D) Cartoon representations of the structure of SARS-CoV-2 RBD in complex with HB27. The RBD is cyan, and the light and heavy chains of HB27 are hotpink and purpleblue, respectively. Residues constituting the HB27 epitope and the RBM are drawn as spheres and colored in green and blue, respectively. The overlapped residues between the HB27 epitope and the RBM are colored in red. The CDRs involved in the interactions with the RBD are labelled. CDR: complementary determining region. RBM: receptor binding motif. (E) Residues in SARS-CoV-2 RBD comprising the HB27 epitope and RBM are labeled. The RBD is rendered as cyan surface. Blue, green and red mark the HB27 epitope, the RBM and overlapped residues of them both, respectively. (F) Hydrophobic interactions between SARS-CoV-2 RBD and HB27. The RBD is shown as cyan ribbon diagrams, and the residues of which involved in hydrophobic interactions with HB27 are shown as side chains and labeled, the four dark orange circles mark the positions of four glycine residues. The HB27 light and heavy chain are rendered as light pink and pale blue molecular surfaces, respectively, of which the residues involved in the hydrophobic interactions with the RBD are highlighted in hotpink and purpleblue and labeled. (G) A few key interactions between SARS-CoV-2 RBD and the HB27 heavy (left) and light chain (right). Hydrogen bonds are presented as dashed lines. See also Figures S3, S4 and S5. Tables S2 and S3.

HB27 binds to the apical head of RBD, partially overlapping with the edge of the RBM core. This binding was independent of glycan recognition (Figure 6D-6E). The head of RBD inserts into the cavity constructed by five complementarity determining regions (CDRs, CDRL1, CDRL3 and CDRH1-3), involving extensive hydrophobic interactions (Figure 6F). The heavy and light chains bury ∼500 Å^2^ and ∼210 Å^2^ of the surface area of the epitope, respectively. Tight binding is further facilitated by 5 hydrogen bonds (Figure 6G and Table S3). HB27 epitope includes 12 residues, of which only 7 residues are conserved between SARS-CoV-2 and SARS-CoV, explaining its specificity for SARS-CoV-2 for binding and neutralization (Figure S5A). Although a number of point mutations in the RBD have been reported in currently circulating strains, none of these mutations lie within the HB27 epitope (Figure S5A). To test the spectrum of neutralizing activities of HB27 against currently circulating strains of SARS-CoV-2, RBD mutants bearing various amino acid substitutions reported were expressed and evaluated for their binding affinities to HB27. In line with structural analysis, all the RBD mutants exhibited comparable binding abilities (Figure S5B). More recently, SARS-CoV-2 isolates encoding a D614G mutation in the C-terminal region of the S1 predominate (Korber et al.). To investigate the neutralizing activities against this more contagious isolate, SARS-CoV-2 PSV harboring the D614G mutation was constructed. Compared to the wild type, HB27 showed similar binding affinities and neutralizing activities against the D614G mutant (Figure S6), indicating that HB27 possibly exhibits broad neutralization activity against SARS-CoV-2 strains currently circulating worldwide.

### Structural dissection of the mechanism of neutralization of SARS-CoV-2 by HB27

Results of our functional studies revealed that HB27 could completely block the interactions of SARS-CoV-2 with ACE2 (Figure 4). To decipher the structural basis for this ability of HB27, the complex structures of SARS-CoV-2 trimer/HB27-Fab and SARS-CoV-2 RBD/ACE2 were superimposed. The superimposition of the structures revealed that HB27 could sterically hinder ACE2 binding (Figure 7A). Out of the 12 residues in the HB27 epitope, 7 residues are involved in tight contacts with ACE2 (Figure 6E and Figure S7). In addition, the three HB27 Fabs act in synergy to abolish ACE2 binding, in which binding of any one ACE2 molecule is sterically hindered by two adjacent HB27 (Figure 7A). Unlike most structural studies of the *apo* SARS-CoV-2 S trimer or complexes with a major configuration corresponding to one ‘open’ RBD and the other two RBDs in ‘closed’ states (Walls et al., 2020; Walls et al., 2019; Wrapp et al., 2020; Zhe Lv, 2020), only one conformational state with one ‘closed’ RBD (mol A) and two ‘open’ RBDs (mol B and C) was observed in our complex structure. Interestingly, Fab-A that binds the closed RBD lies between two open RBDs, forming contacts with the mol B-RBD and the Fab-C located in proximity to the mol C-RBD (Figure 7B). Probably acting as a bridge, the Fab-A, to some extent, anchors links of all three RBDs and restrains their conformational transitions (Figure 7B). Perhaps correlated with this, HB27 possesses the ability to disrupt the membrane fusion event through restraining the conformational changes playing out during the progression from the prefusion to the postfusion state. Collectively these data suggest that HB27 might prevent both the attachment of SARS-CoV-2 to host cells and viral fusion with endosomal membrane. However, fusion blockade by HB27 might be dependent on the uptakes of antibodies into the endosome.

**Figure 7.**
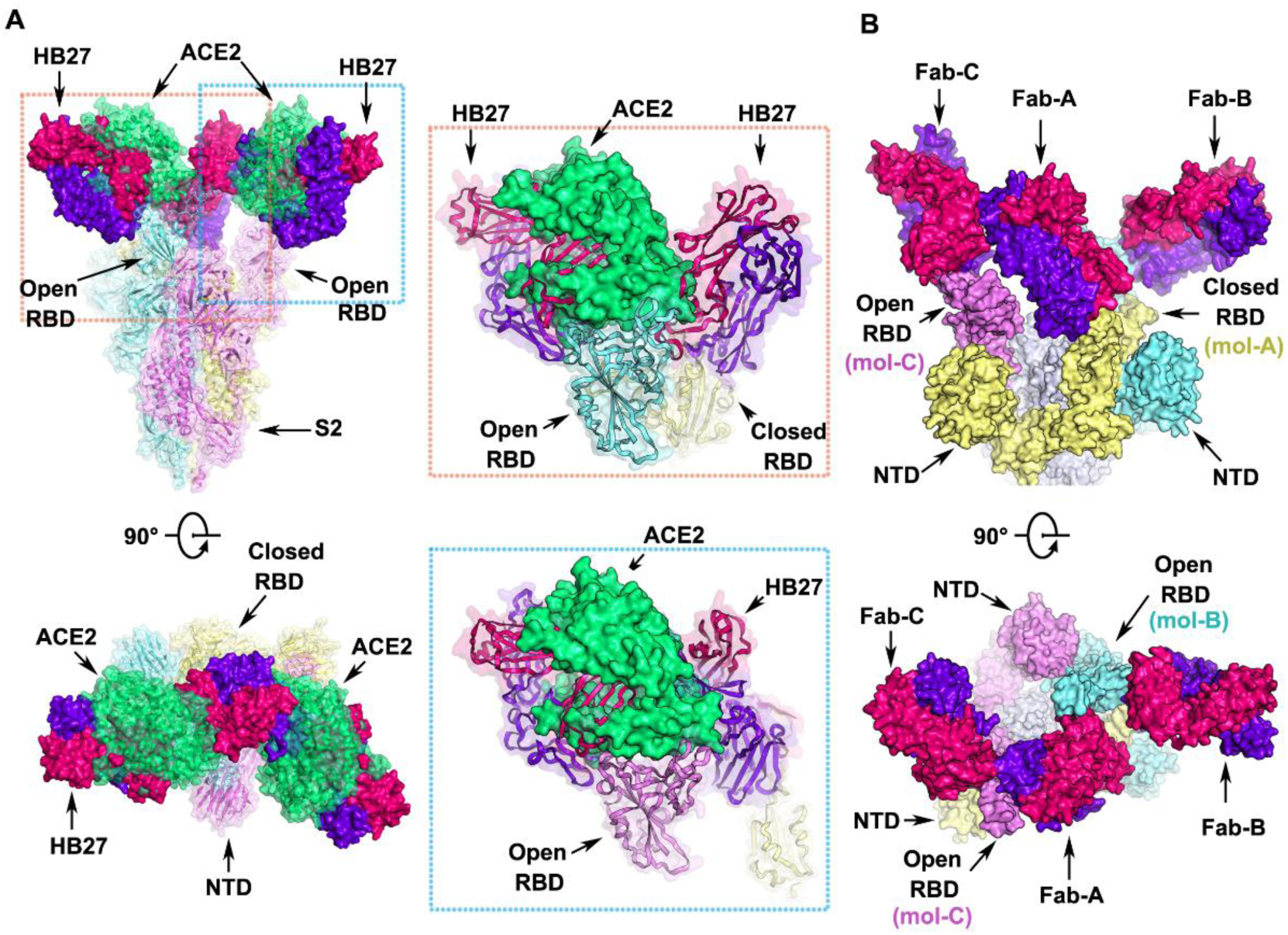
Structural basis for neutralization of SARS-CoV-2 by HB27. (A) Orthogonal views of the clashes between HB27 Fabs and ACE2 upon binding to SARS-CoV-2 S trimer. The SARS-CoV-2 S trimer is presented as ribbon diagrams and translucent molecular surfaces with three monomers colored in cyan, yellow and violet, respectively. The three copies of HB27 Fabs are rendered as molecular surfaces colored the same as in Figure 6. The superposed ACE2 is presented as green ribbon diagrams as well as translucent molecular surface. Insets are close-up views of the clashes between ACE2 and HB27 upon binding to SARS-CoV-2 RBD. (B) Orthogonal views of the structure of HB27 Fab-A, Fab-B and Fab-C complexed with SARS-CoV-2 RBD. The S1 subunits of SARS-CoV-2 S trimer are rendered as cyan, yellow and violet surfaces and the S2 subunits are rendered as gray surfaces. See also Figures S6 and S7.

## Discussion

SARS-CoV-2 shares about 80% sequence identity with SARS-CoV, implying that both these viral strains share a similar mechanism of establishing an infection, including targeting a similar spectrum of host cells, employing a similar entry pathway and hijacking the same cellular receptor (Hoffmann et al., 2020; Zhou et al., 2020). Both cross-reactive and virus-specific human NAbs have been identified, despite around 77% of amino-acid sequence identity between the S of SARS-CoV-2 and SARS-CoV (Brouwer et al., 2020; Hansen et al., 2020; Pinto et al., 2020; Wec et al., 2020; Wu et al., 2020; Yuan et al., 2020; Zhe Lv, 2020). It is important to decipher the immunogenic mechanism to discover patterns of different patches comprising different residues eliciting cross-reactive or virus-specific NAbs with various neutralization mechanisms. Currently, several cross-reactive mAbs, including CR3022, H014 and S309, screened from convalescent SARS patients or *via* immunization using SARS-CoV RBD, show distinct neutralizing activities against SARS-CoV-2 (Pinto et al., 2020; Yuan et al., 2020; Zhe Lv, 2020). Structural analysis reveals that all these mAbs recognize conserved patches either distal from or proximal to the edge of the RBM, but not in the RBM. Interestingly, the corresponding epitope in both open and closed RBDs is accessible to S309, but accessible to H014 only in open RBDs, and can only be accessed by CR3022 when at least two RBDs are in the open conformation. The stoichiometric binding of Fab to the S trimer might correlate with the neutralizing activities, probably explaining the weak neutralization efficiency observed for CR3022. HB27 targets the less conserved edge of the RBM core with a full occupancy for all RBDs. This structural observation supports the observed specificity of HB27 for SARS-CoV-2 and its highly potent neutralization of SARS-CoV-2.

Our results indicate that HB27 probably inhibits SARS-CoV-2 infection at multiple steps during the viral entry process. First, viral infection can be stalled by hindering the attachment of SARS-CoV-2 to host cells by preventing interactions between the RBD and ACE2, which is the major neutralization mechanism for most RBD-targeting NAbs. Upon virus attachment and entry into host cells, proteolytic activation at the S1/S2 boundary leads to S1 dissociation and a dramatic structural change in S2, which triggers viral membrane fusion (Shang et al., 2020). To date, antibodies that are capable of interfering with coronavirus fusion have not been reported. HB27 may be involved in restraining the conformational changes required for the progression of the life cycle of the virus from the prefusion to the postfusion stage. Furthermore, recent studies suggest that the SARS-CoV-2 entry depends on ACE2 and cell surface protease TMPRSS2 (Hoffmann et al., 2020; Ou et al., 2020). A blockage of viral attachment to host cell surface by HB27 possibly affects the colocalization of SARS-CoV-2 S with TMPRSS2 on the cell membrane. This may be yet another way employed by HB27 to prevent viral membrane fusion where the cleavage of S by TMPRSS2 is averted. Therefore, the potent neutralizing activity of HB27 probably results from its intervention at two steps of viral infection, locking away attachment of the virus to its receptor and blocking membrane fusion; resulting in a double lock.

Most importantly, the *in vivo* protection efficacy of HB27 was confirmed in two established mouse models. The results of these studies consistently showed that a single dose of HB27 either before or post SARS-CoV-2 exposure not only blocked viral replication in the lungs and trachea, but also prevented the pulmonary pathological damage. To date, only a few neutralizing antibodies have been tested in animal models (Cao et al., 2020; Shi et al., 2020). Previously, we have shown that H014 reduced pulmonary viral loads by ∼100-fold in human ACE2 mice (Zhe Lv, 2020). HB27 exhibits a more potent protective efficacy in reducing viral RNAs (∼11,000-fold) with a much lower administration dose (20 mg/kg v.s. 50 mg/kg). The preliminary results on the efficacy of the antibody as well as the safety profile of HB27 in Rhesus macaques support testing of its potential in curing COVID-19 in clinical trials. In fact, while this manuscript was under review, HB27 entered clinical trials in China (registration number NCT04483375). More details can be found at *https://clinicaltrials.gov/ct2/show/NCT04483375?cond=SCTA01&draw=2&rank=1*. bIn summary, our results not only show how increasing access to panels of authentic neutralizing monoclonal antibodies will facilitate structure-function studies to unpick the underlying biological processes of virus-host interactions, but also provide molecular basis for applying HB27 for potential COVID-19 treatment, highlighting the promise of antibody-based therapeutic interventions.

## Acknowledgements

We thank Dr. Xiaojun Huang, Dr. Boling Zhu, Xianjin Ou and Dr. Gang Ji for cryo-EM data collection, the Center for Biological imaging (CBI) in Institute of Biophysics for EM work. Work was supported by the National Key Research and Development Program (2020YFA0707500, 2018YFA0900801, 2017YFC0840300), the Strategic Priority Research Program (XDB29010000, XDB37030000), National Science and Technology Major Projects of Infectious Disease funds (grants 2017ZX103304402), National Natural Science Foundation of China (NSFC) (grants 82041005, 12034006) and the Beijing Municipal Science and Technology Project (Z201100005420017). Xiangxi Wang was supported by Ten Thousand Talent Program and the NSFS Innovative Research Group (No. 81921005). Chengfeng Qin was supported by the National Science Fund for Distinguished Young Scholar (No. 81925025), and the Innovative Research Group (No. 81621005) from the NSFC, and the Innovation Fund for Medical Sciences (No.2019-I2M-5-049) from the Chinese Academy of Medical Sciences. Ling Zhu was supported by the Youth Innovation Promotion Association at the Chinese Academy of Sciences (2019098).

## Author contributions

X.W., C.F.Q., L.Z., Z.R. and C.S. conceived, designed and supervised the study and prepared this manuscript. X.W., C.F.Q., L.X. and Y.W. coordinate the project. Z.C., Zhe.L. and Y.S. purified proteins; Y.D., R.Z., Q.C., N.Z., Q.Y., X.L. and T.C. performed live virus and animal assays; C.S., H.W., D.K., J.M., C.L. Y.Z. and L.X. generated antibodies, constructed mutants and carried out safety evaluations in macaques; L.C., Z.C., and Y.S prepared cryo-EM grids and collected cryo-EM data; N.W., L.W., Z.C. and X.W. processed data; L.Z. and X.W. built and refined the structure model; L.Z., N.W. and X.W. analyzed the structures; X.X. performed liposome membrane fusion assay; C.F., W.H., J.N., Q.L. and Y.W constructed PSV, PSV-related mutants and PSV-based neutralization. All authors discussed the experiments and results, read, and approved the manuscript.

## Data and materials availability

Cryo-EM density maps have been deposited at the Electron Microscopy Data Bank with accession codes EMD-30503 (complex) and EMD-30500 (binding interface) and related atomic models has been deposited in the protein data bank under accession code 7CYP and 7CYH, respectively.

## STAR Methods

### Facility and ethics

Experiments involving live SARS-CoV-2 virus were performed in the enhanced biosafety level 3 (P3+) facilities in the Institute of Microbiology and Epidemiology, Academy of Military Medical Sciences. All animal experiments were approved by the Experimental Animal Committee of Laboratory Animal Center, AMMS (approval number: IACUC-DWZX-2020-001).

### Cells and viruses

The human embryonic kidney 293T cell line (Cat: CRL-11268) used for pseudovirus (PSV) packaging was purchased from ATCC. Vero-E6 cells were purchased from Chinese Academy of Medical Sciences Cell Bank (Cat: GN017). Vero cells, 293T and Vero-E6 cells were grown in Dulbecco’s modified Eagle’s medium (DMEM) containing 10% (v/v) FBS. The SARS-CoV-2 viral strain BetaCoV/Beijing/IME-BJ01/2020 was originally isolated from a COVID-19 patient returning from Wuhan, China. The virus was amplified and titrated by standard plaque forming assay using Vero cells.

### Protein expression and purification

Plasmids for protein expression were constructed by inserting the genomic sequences of SARS-CoV RBD (residues 306–527, GenBank: NC_004718.3), SARS-CoV-2 RBD (residues 319-541, GenBank: MN908947.3), and SARS-CoV-2 S trimer (residues 1– 1208, GenBank:MN908947.3), respectively, into the mammalian expression vector pCAGGS with a C-terminal 2×StrepTag. Proline substitutions at residues 986 and 987, a “GSAS” instead of “RRAR” at the furin cleavage site were performed on the gene encoding S protein based on the research of Jason S. McLellan (Wrapp et al., 2020). Polyethylenimine was used to transiently transfect HEK Expi 293F cells (Thermo Fisher) with SARS-CoV RBD, SARS-CoV-2 RBD and SARS-CoV-2 S, respectively. StrepTactin resin (IBA) was used for protein purification from the cell supernatants, followed by size-exclusion chromatography with a Superose 6 10/300 column (GE Healthcare) or a Superdex 200 10/300 Increase column (GE Healthcare) in 20mM Tris, 200 mM NaCl, pH 8.0.

### Reagents, recombinant proteins and antibodies

Recombinant RBD protein of SARS-CoV-2 with His tag (Cat: 40150-V08B2, monomer in solution), Recombinant ACE2 protein with His tag (Cat: 10108-H08H, monomer in solution), transfection reagent Sinofection (Cat: STF02), mammalian expression plasmids of full-length S protein with GFP tag at the C terminal (Cat: VG40589-ACGLN) were purchased from Sino Biological. Fetal bovine serum (FBS) (Cat: SA 112.02) were purchased from Lanzhou Minhai Bio-engineering. Luciferase assay system (Cat: E1501) was purchased from Promega. Anti-human IgG Fc/HRP (Cat: 5210-0165) were purchased from KPL. Goat anti-human IgG F(ab’)2/HRP (Cat: 109-036-006) were purchased from Jackson ImmunoResearch.

### Generation of humanized anti-SARS-CoV-2 antibody HB27

SARS-CoV-2 antibodies were screened from a phage-display scFv library constructed from the spleen mRNA of mice immunized with recombinant SARS-CoV-2 RBD protein. SARS-CoV-2 RBD was used as the bait to select for specific anti-RBD scFvs by biopanning and the scFvs exhibiting potent binding for SARS-CoV-2 RBD were generated as chimeric antibodies. The chimeric antibodies were expressed using HEK-293T transient transfection production system and examined for competition activities with ACE2 for binding to SARS-CoV-2 RBD and neutralizing activities against SARS-CoV-2 and SARS-CoV pseudoviruses. The chimeric antibody mhB27 exhibited high binding affinity to SARS-CoV-2 RBD and potent neutralizing activity against SARS-CoV-2 pseudoviruses, therefore its humanized version-HB27 (Fc modified IgG1 subtype) was further generated.

### Generation of Fab fragment

The HB27 Fab fragment was prepared using Pierce FAB preparation Kit (Thermo Scientific) following the manufacturer’s instructions. In brief, following removal of the salt with a desalting column, the antibody was mixed with papain and incubated for digestion at 37 °C for 3-4 h. The HB27 Fab was separated using protein A affinity column and concentrated for further applications.

### Generation of mutant RBDs

Genomic information of SARS-CoV-2 mutant strains were obtained from GISAID (https://platform.gisaid.org), selected site mutants within the RBD domain (residues 306-527) were conducted. The mutated RBD genes with His-tag were cloned into pSTEP2 vector and transfected into 293T cells for protein expression. Cell culture supernatants were collected and purified using IMAC resins.

### Protein-protein interaction identified by Octet

Recombinant SARS-CoV-2 RBD-His was biotinylated and loaded onto SA sensor (Pall corporation), and then HB27 antibody or HB27 Fab fragments were added for real-time association and dissociation analysis using Octet96e (Fortebio). Data was processed with Data Analysis Octet.

### ELISA

The competition between HB27 and ACE2 for binding to SARS-CoV-2 RBD, and the binding of HB27 antibody to mutant SARS-CoV-2 RBDs are examined by ELISA. Recombinant RBD protein was coated on 96-well plates using CBS buffer over night at 4°C. The plates were blocked in BSA at room temperature for 1 h. Recombinant ACE2 with an His-tag and serial diluted HB27 antibody were then added and incubated at room temperature for 1 h. After washing away the unbound proteins and antibodies, secondary antibody against His-tag with HRP labeling were added and incubated for 1 h before washed away. Developing buffer was added and incubated for 5-30 min, 1% H_2_SO_4_ was added to stop the reaction and absorbance at 450 nm was detected with a microplate reader. Recombinant RBD mutant proteins were coated on 96-well plates using PBS buffer at 4 °C for 12 h. After that BSA solution was used for blocking at 25 °C for 1 h. Serial diluted antibodies were then added and incubated at room temperature for 1 h. After washing away the unbound antibodies, secondary antibody against human IgG with HRP labeled was added and incubated for 1 h before washed away. For color development, TMB mixture solution was added and incubated for 5-30 min, then 1% H_2_SO_4_ was added to stop the reaction and absorbance at 450 nm was detected with a microplate reader.

### Flow cytometry

HB27 was serial diluted and incubated with 293T-ACE2 cells or 293T-SARS-CoV-2-S cells together with recombinant SARS-CoV-2 RBD or ACE2 for 45 min, respectively. Following the washing away of unbound proteins, cells were incubated with FITC labeled secondary antibody for 20 min and subject to flow cytometer for examination of cellular binding. Data were analyzed using Flowjo and Graphpad.

### Production of pseudoviruses

Pseudoviruses were prepared as previously described (Nie et al., 2020). In brief, 293T cells were transfected with the plasmids of SARS-CoV S or SARS-CoV-2 S, respectively. 24 hours later, transfected 293T cells were infected with VSV G pseudotyped virus (G*ΔG-VSV) at a multiplicity of infection (MOI) of 4. Two hours post infection, cells were washed three times using PBS, followed by adding complete culture medium. Twenty-four hours post infection, SARS-CoV or SARS-CoV-2 pseudoviruses were harvested, 0.45-μm filtered and stored at −80 °C.

### Pseudovirus neutralization assay

Aliquots of a 100 μL of ∼40,000 Vero-E6 cells/well were added into 96-well plates. 60 μL of SARS-CoV/SARS-CoV-2 pseudoviruses and 60 μL of serial diluted antibody samples were incubated for 1 h at 37°C, after which the pseudovirus-mAb mixtures were added into the wells containing Vero-E6 cells. The 96-well plates were then incubated for 24 hours in a 5% CO_2_ environment at 37°C, then the luciferase luminescence (RLU) was measured using luciferase assay system following the manufacturer’s manual with a luminescence microplate reader. The neutralization percentage was calculated by the formula: Inhibition (%) = [1-(sample RLU-Blank RLU)/(Positive Control RLU-Blank RLU)] (%). Neutralization titers of the antibodies were presented as 50% maximal inhibitory concentration (IC_50_).

### Immunofluorescence

293T cells were transfected with SARS-CoV-2-S-GFP or ACE2-GFP. 48h later, cells were fixed with 4% paraformaldehyde (PFA) for 20 min at room temperature and stained for nuclei with 4,6-diamidino-2-phenylindole (DAPI). HB27 antibody was incubated for 1h, followed by incubation of RBD-His and anti-His-PE,or APC labelled ACE2-Fc for 20 min. The fluorescence images were recorded using a Nikon A1 confocal microscope.

### Liposome preparation

Lipids, 1-palmitoyl-2-oleoyl-sn-glycero-3-phosphocholine (POPC; Avanti-Polar Lipids), 1,2-dioleoyl-sn-glycero-3-phospho-L-serine (DOPS; Avanti-Polar Lipids), 1,2-dihexadecanoyl-sn-glycero-3-phosphoethanolamine (Texas Red-DHPE; Sigma ChemicalCo.) were mixed in a 84.5:15:0.5 molar ratio and prepared as previously reported (Qiu et al., 2018). The dried lipid film was hydrated at room temperature with 100 mM calcein (Sigma) in buffer (25 mM HEPES, 150 mM KCl, pH 7.4), and then the vesicles were extruded 25 times using the Mini-Extruder device (Avanti Polar Lipids) through Nuclepore filters (Whatman) with a pore size of 0.1 μm. Unincorporated calcein was separated from the liposomes using a Sephadex G-50 column. Liposomes (10 mM lipid on the basis of the input lipid) were stored at 4°C and used within 1 week.

### Liposome-binding and Calcein-leakage assays

SARS-CoV-2 (∼20 μg) was incubated with 0.1 μM trypsin (Sigma) at 37°C for 20 min. Then the virus was mixed with 0.3 μM ACE2 and HB27 antibody with the final concentration of 2.5 μM, 0.5 μM, 0.1 μM or 0.02 μM. The mixture was added to 0.1 mM liposomes in a total volume of 90 μl in a 96-well plate, and the fluorescence (excitation at 460 nm, emission at 509 nm) was monitored at 37 °C using a SpectraMax M5 Microplate Reader (Molecular Devices). At t = 0 sec, the pH of the medium were adjusted to 5.6 by addition of 10 μl of 1 M MES (morpholineethanesulfonic acid, pH 5.6) as *F*_0_. The emission fluorescence was recorded as *F*_t_ at 10 sec intervals. After 500 sec, 10 μl of 10% Triton X-100 was added to achieve complete release of the maximum fluorescence as *F*_100_. The fusion scale was calibrated such that 0% fusion corresponded to the initial excimer fluorescence value. The percentage of calcein leakage at each time point is defined as: leakage (%) = (*F*_t_ - *F*_0_) × 100 / (*F*_100_ - *F*_0_).

### Cell–cell fusion assay

The establishment and detection of cell–cell fusion assay was performed as previously described (Xia et al., 2020). In brief, Vero-E6 cells were used as target cells and 293T cells transfected with SARS-CoV-2 S-GFP protein expression vectors were served as effector cells. Effector cells and target cells were co-cultured in the absence or presence of antibodies in DMEM containing 10% FBS for 48 h. After incubation, cells were fixed with 4% paraformaldehyde (PFA) at room temperature for 20 min and stained for nuclei with 4,6-diamidino-2-phenylindole (DAPI). The fluorescence images were recorded using a Leica SpeII confocal microscope. S-mediated cell-cell fusion was observed by the formation of multi-nucleated syncytia. Five fields were automatically collected in each well to count the number of fused and unfused cells and the antibody inhibition rate was calculated as following: fusion rate (FR) = (fused cell number) / (fused cell number+ unfused cell number), Inhibition%= (Positive Control (FR) – Sample (FR)) / (Positive Control (FR)) %. The experiment was performed three times.

### Negative stain

Samples were diluted to a desired concentration (∼0.02 mg/mL) and deposited onto freshly glow-discharged carbon-coated grids. After rinsing twice with deionized water, the grids were stained with 1% phosphotungstic acid (pH 7.0) and loaded onto a 120-kV transmission electron microscope (FEI) for inspection.

### Cryo-EM sample preparation and data collection

HB27 Fab fragments and SARS-CoV-2 S ectodomain (1mg/ml) were purified and incubated at a ratio of 9 Fab molecules per S trimer. 3μL aliquots of the mixture were applied onto freshly glow-discharged C-flat R1.2/1.3 Cu grids. The grids were blotted for 3 s in 100% relative humidity for plunge-freezing (Vitrobot; FEI) in liquid ethane. The Cryo-EM data sets were collected at 300 kV with a Titan Krios microscope (Thermo Fisher) fitted with a Gatan K2 detector. Movies (32 frames, each 0.2 s, total dose 60 e^−^Å^−2^) were recorded at defocuses of between 1.25 and 2.7 μm using SerialEM, yielding a final pixel size of 1.05 Å.

### Image processing

Micrographs of SARS-CoV-2 S trimer-HB27 Fab complex were recorded. The defocus values for each micrograph was determined using Gctf (Zhang, 2016). Then particles were picked and extracted for 2D alignment and 3D classification by using the *apo* structure of SARS-CoV-2 S trimer (Walls et al., 2020) as the initial model in Relion (Scheres, 2016). The best classes were selected and used for 3D refinement and postprocessing (estimate the B-factor automatically), yielding the final resolution of 3.5 Å based on the gold-standard Fourier shell correlation (threshold = 0.143) (Scheres and Chen, 2012). However, the densities for the binding interface between RBD and HB27 are weak due to the conformational heterogeneity of the RBD. To solve this problems, we utilized the block-based reconstruction strategy (Wang et al., 2020; Wang et al., 2019; Yang et al., 2020) for focusing classification and refinement. Details on parameter settings can be found in structural determinations for the binding interface between RBD and H014 (Zhe Lv, 2020). In addition, local averaging of the RBD-Fab equivalent copies present in different classes further improves the resolution to 3.9 Å. All procedures were performed with Relion (Scheres, 2016). The local resolution was evaluated by ResMap (Kucukelbir et al., 2014).

### Model building and refinement

The structures of SARS-CoV-2 S trimer and a human Fab fragment (Protein Data Bank ID: 6VSB and 5N4J, respectively) were manually fitted into the refined map of SARS-CoV-2 S trimer-HB27 complex in Chimera (Pettersen et al., 2004) and then improved by manual real-space refinement in COOT (Brown et al., 2015). The atomic model was further subject to real-space positional and B-factor refinement using Phenix (Afonine et al., 2012). The final models were evaluated using Molprobity (Chen et al., 2010). Detailed informatin of the data sets and refinement statistics are summarized in Table S2.

### Surface plasmon resonance

The SARS-CoV-2 S trimer was immobilized onto a CM5 sensor to ∼500 response units (RUs) using Biacore 8K (GE Healthcare). Serial diluted HB27 or Fab fragments or recombinant ACE2 flowed through the sensor. For competitive binding assays, the first sample was allowed to flow over the chip at a rate of 20 μl/min for 120 s, and then the second sample was injected at the same rate for another 120 s. The response units were recorded and analyzed.

### Plaque reduction neutralization tests (PRNT)

The neutralization activity of HB27 against SARS-CoV-2 were examined by standard plaque reduction neutralization tests (PRNT) in Vero cells. In brief, 5-fold serial dilutions of HB27 were mixed with ∼100 PFU of SARS-CoV-2 and incubated at 37 °C for 1 hour. The mixture was then added to Vero-E6 cell monolayers in a 12-well plate in duplicate and incubated at 37 °C for 1 hour. After which the mixture was removed, and 1 ml of 1.0% (w/v) LMP agarose (Promega) in DMEM supplemented with 4% (v/v) FBS was layered onto the infected cells. Following a two-day incubation at 37 °C, the wells were stained with 1% (w/v) crystal violet in 4% (v/v) formaldehyde for plaque visualization. The PRNT_50_ values were determined using non-linear regression analysis with GraphPad prism.

### Protection against SARS-CoV-2 challenge in hACE2 mice

The *in vivo* protection efficacy of HB27 antibody was evaluated using a newly established mouse model based on a SARS-CoV-2 mouse adapted strain MASCp6 (Gu et al., 2020) and a humanized hACE2 mouse model (Sun et al., 2020), respectively. Briefly, a group of 6 to 8-week-old hACE2 humanized mice or BALB/c mice were intraperitoneally administrated with HB27 (20 mg/kg) before (prophylactic) and/or after (therapeutic) challenge with 5 × 10^5^ PFU of SARS-CoV-2 or 1.6×10^4^ PFU of MASCp6 via intranasal route, respectively. All mice were monitored daily for morbidity and mortality. The lung tissues of mice were collected at 3 and 5 dpi for viral RNA loads assay and HE staining.

### Viral RNA quantitation

Viral RNA quantification was performed by RT-qPCR aplying One Step PrimeScript RT-PCR Kit (Takara, Japan). The primers and probe targeting against the gene of SARS-CoV-2 S used for RT-qPCR were CoV-F3 (5’-TCCTGGTGATTCTT CTTCAGGT-3’); CoV-R3 (5’-TCTGAGAGAGGGTCAAGTGC-3’); and CoV-P3 (5’-FAM-AGCTGCAGCACCAGCTGTCCA -BHQ1-3’), respectively.

### Pre- and post-adsorption inhibition assay

Pre- and post-adsorption inhibition assays were performed as described previously (Wang et al., 2017). For the post-adsorption assay, SARS-CoV-2 was first added to Vero cells for 1 hour at 4 °C, and then the cells were washed three times, following which the mAb was added and incubated for 1 hour at 4 °C. For the pre-adsorption assay, the mAb was firstly incubated with SARS-CoV-2 for 1 hour at 4 °C before the mAb-virus mixture was added to Vero cells. After three washes using PBS, the PRNT was performed as described above. And the detection of the remaining amount of SARS-CoV-2 RNA on the surface of Vero cells after HB27 treatment was carried out with quantitative RT-PCR.

### Histology and Immunostaining

Mouse tissues were excised and fixed with 10% neutral buffered formaline, and then dehydrated and embedded in paraffin. Sections of 4 μm thickness were obtained and stained with hematoxylin and eosin (H & E) following standard histological procedures. Images were recorded using Olympus BX51 microscope equipped with a DP72 camera.

### Toxicokinetics of HB27 in Rhesus Monkeys

Rhesus macaques were randomly grouped into two groups, one group was given placebo and one group was given a single dose of HB27 at 500 mg/kg intravenously. Blood samples were collected at pre-dose, immediately after completion of dosing (± 1 minute), and 1h, 2h, 4 h, 8 h, 24 h (Day 2), 48 h (Day 3), 72(Day4), 120 h (Day 6), 168 h (Day 8) and 336 h (Day 15) after beginning of infusion. Serum concentration of HB27 was measured using ELISA.

### Clinical pathology of HB27 in Rhesus Monkeys

Blood samples were collected via forelimb or hindlimb subcutaneous vein at predose and 3, 8 and 16 days postdose. Hematology parameters including white blood cells (WBC), lymphocytes (Lymph), red blood cells (RBC) and hemoglobin concentration (HGB) were measured using an ADVIA Hematology system. Clinical chemistry parameters including AST (aspartate transaminase), ALT (alanine transaminase) and ALP (alkaline phosphatase) and LDH (lactate dehydrogenase) were measured using TBA-120FR. BD FACS Calibur Flow Cytometry was used for determinations of CD4+, CD8+ T percentages.

## Supplemental Information for

**Figure S1.**
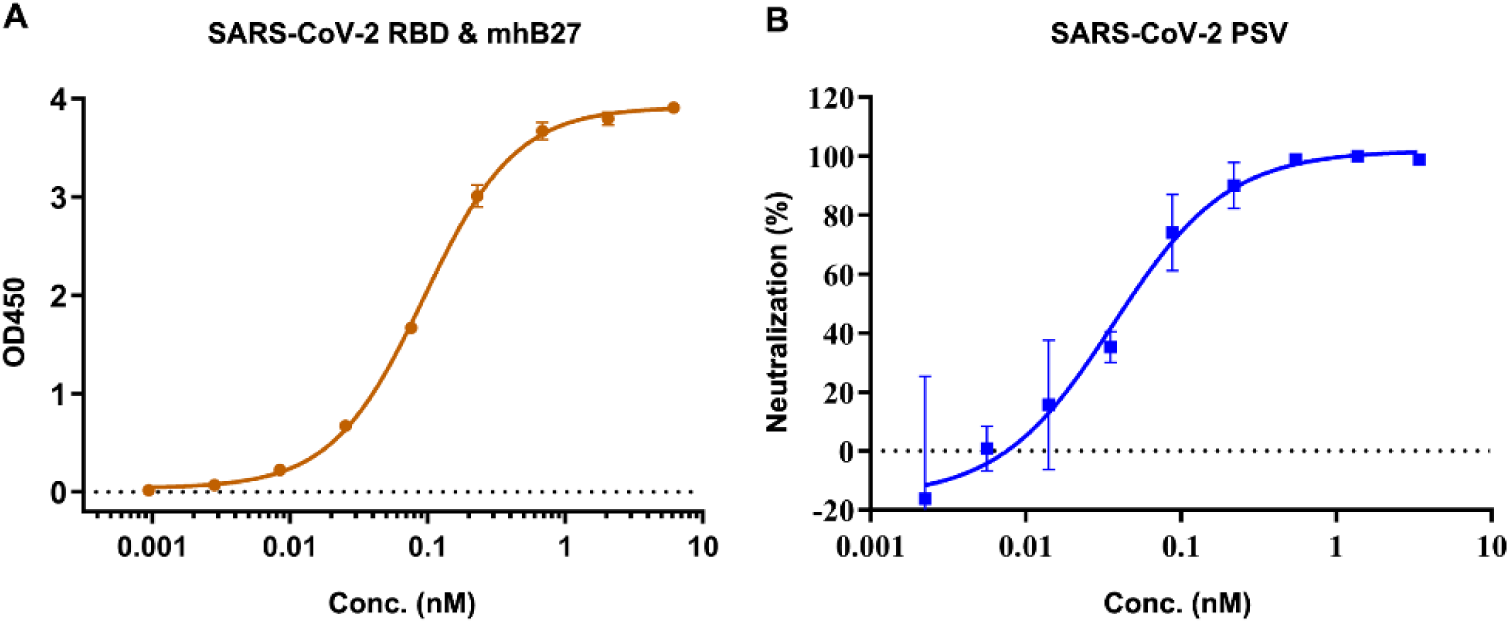
Murine antibody mhB27 strongly binds SARS-CoV-2 RBD and neutralizes SARS-CoV-2 PSV. Related to Figure 1. (A) Binding assay of mhB27 to SARS-CoV-2 RBD. mhB27 was serial diluted and tested its ability to bind to SARS-CoV-2 RBD by ELISA. (B) Neutralizing activities of mhB27 against SARS-CoV-2 pseudoviruses (PSV).

**Figure S2.**
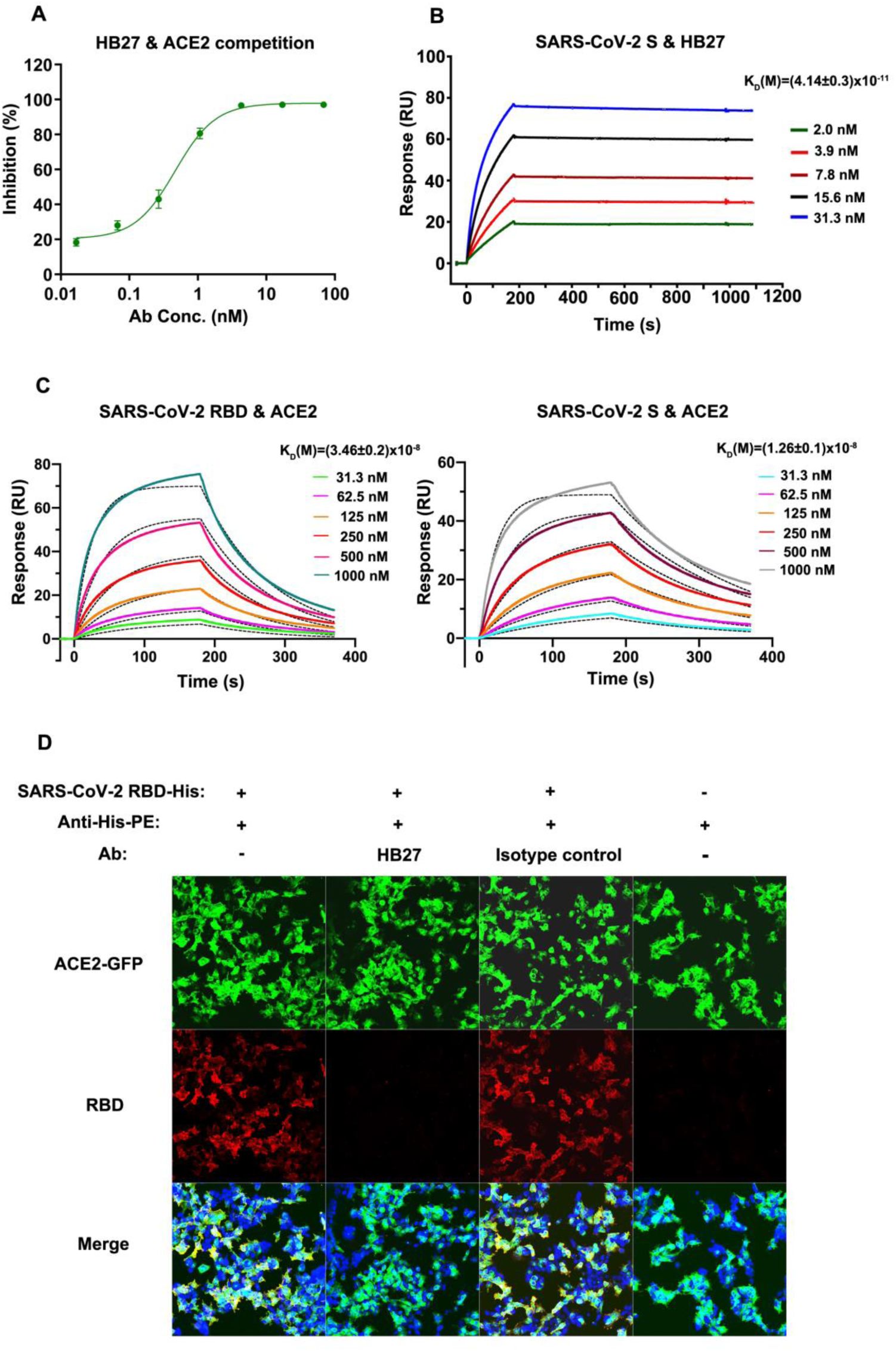
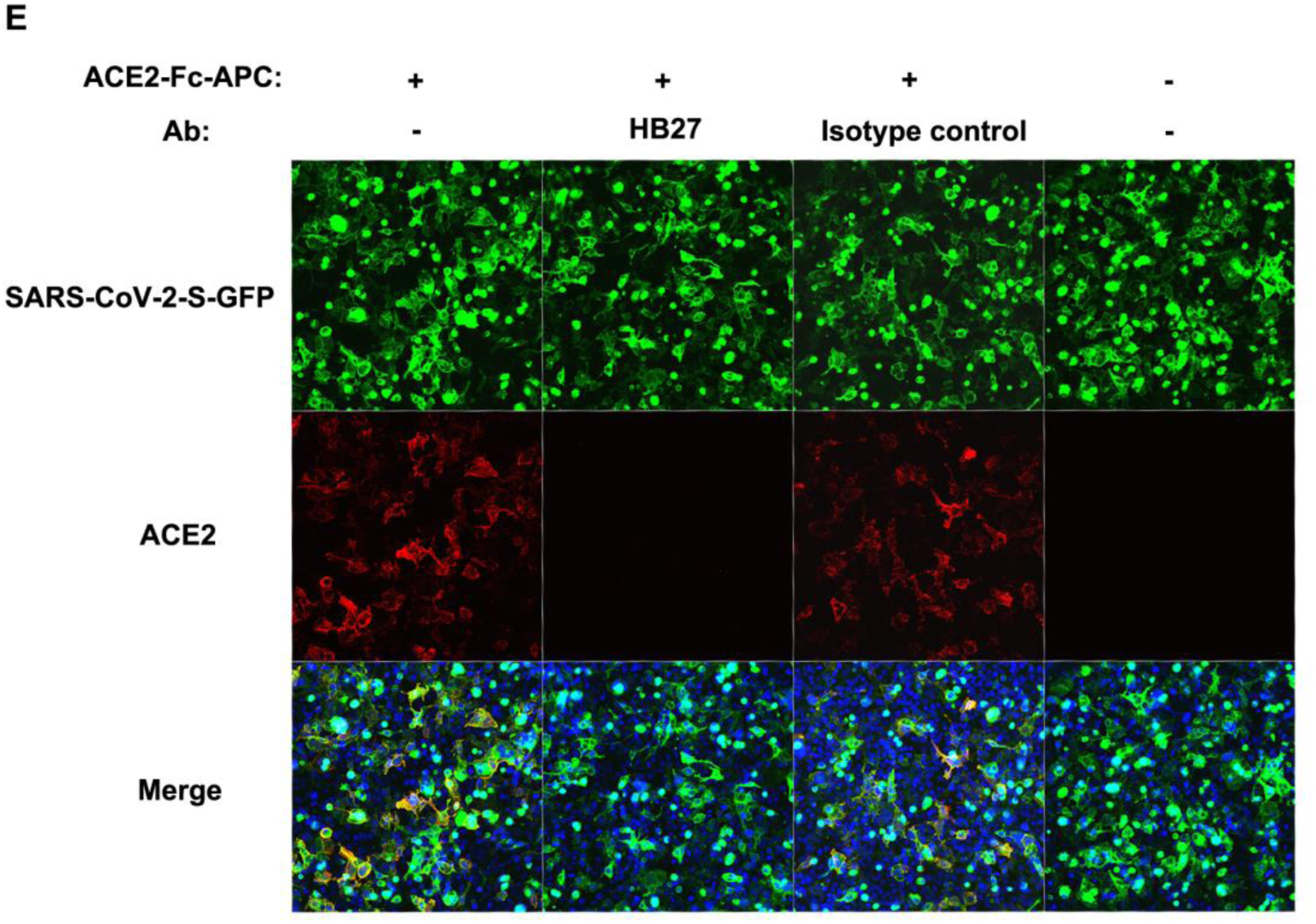
HB27 potently competes with ACE2 for binding to SARS-CoV-2 RBD. Related to Figure 4. (A) HB27 was demonstrated to compete with recombinant ACE2 for binding to SARS-CoV-2 RBD with an EC_50_ value of 0.5 nM by the enzyme-linked immunosorbent assay (ELISA). (B) BIAcore SPR kinetic profile of SARS-CoV-2 S trimer and HB27. The binding affinity K_D_ (equilibrium dissociation constant, K_D_ = Kd/Ka, where Kd and Ka represent the dissociation rate constant and association rate constant, respectively) values were obtained using a series of HB27 concentrations and fitted in a global mode in each sensorgram. (C) BIAcore SPR kinetic profiles of SARS-CoV-2 RBD (left panel) and S trimer (right panel) with ACE2. The binding affinity K_D_ (equilibrium dissociation constant, K_D_ = Kd/Ka, where Kd and Ka represent the dissociation rate constant and association rate constant, respectively) values were obtained using a series of HB27 concentrations and fitted in a global mode in each sensorgram. (D) Competition of HB27 for SARS-CoV-2 RBD binding to 293T cells expressing GFP-tagged ACE2 as detected by immunofluorescence assay, scale bar, 100 μm. Anti-H7N9 mAb was used as an isotype control. (E) Competition of HB27 for ACE2-Fc-Apc binding to 293T cells expressing GFP-tagged SARS-CoV-2-Spike as detected by immunofluorescence assay. Anti-H7N9 mAb was used as an isotype control.

**Figure S3.**
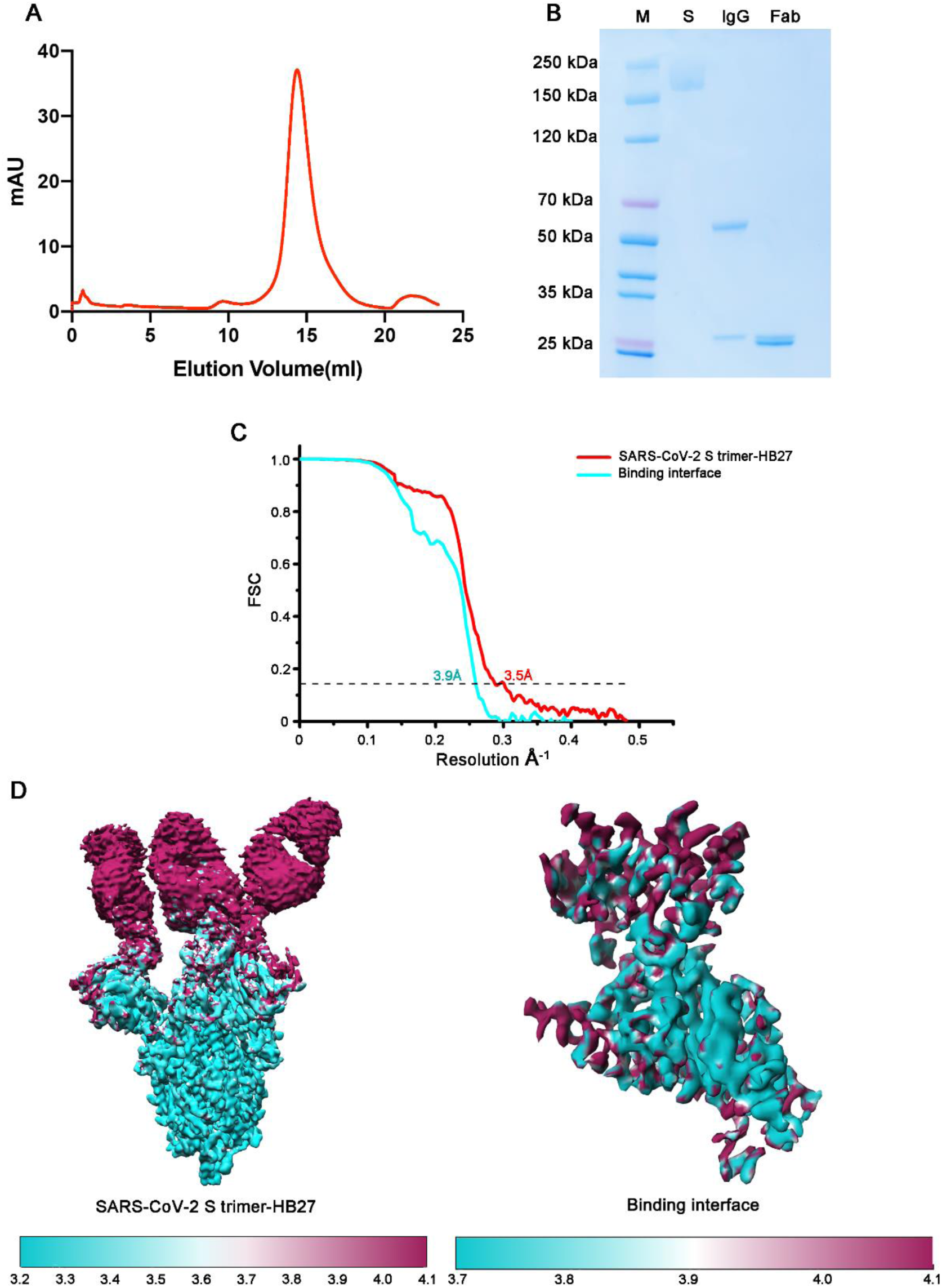
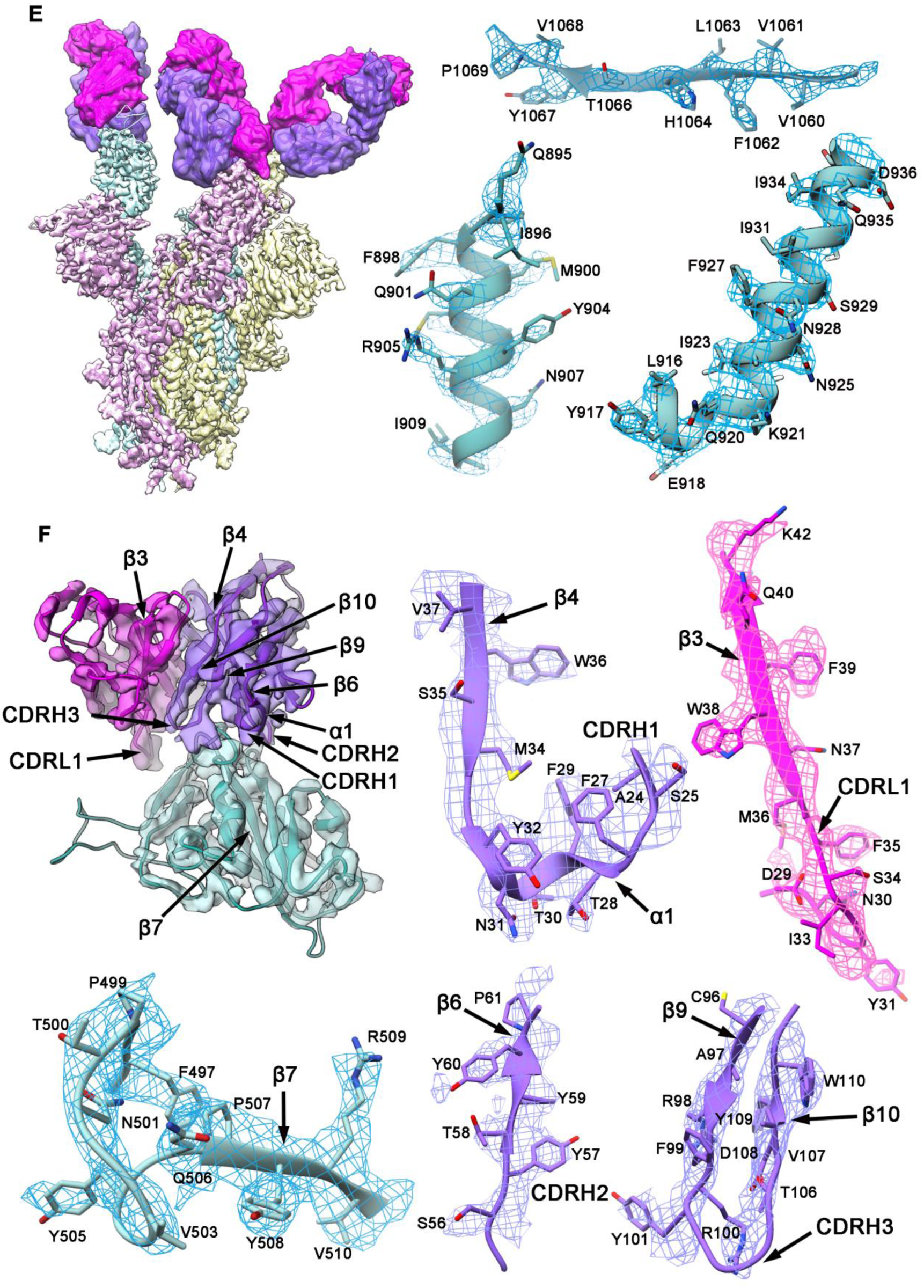
Characterization of SARS-CoV-2 and HB27, and cryo-EM maps and atomic models of SARS-CoV-2 S and HB27 complex. Related to Figure 6. (A) Gel filtration of SARS-CoV-2 S trimer. (B) SDS-PAGE analysis of the SARS-CoV-2 S trimer, the HB27 IgG and the Fab fragment. (C) The gold-standard Fourier Shell Correlation (FSC) curves of the final cryo-EM maps of the SARS-CoV-2 S trimer-HB27 Fabs complex and of the binding interface. (D) Local resolution evaluations of the cryo-EM maps of SARS-CoV-2 S trimer complexed with three HB27 Fabs and the binding interface using ResMap (Kucukelbir et al., 2014) are shown. (E) Cryo-EM map of SARS-CoV-2 S trimer complexed with three HB27 Fabs. (F) Cryo-EM map of the binding interface between SARS-CoV-2 RBD and one HB27 Fab. The color scheme is the same as in Figure 6. The magnified panels illustrate both maps (mesh) and related atomic models. Residues are shown as sticks,

**Figure S4.**
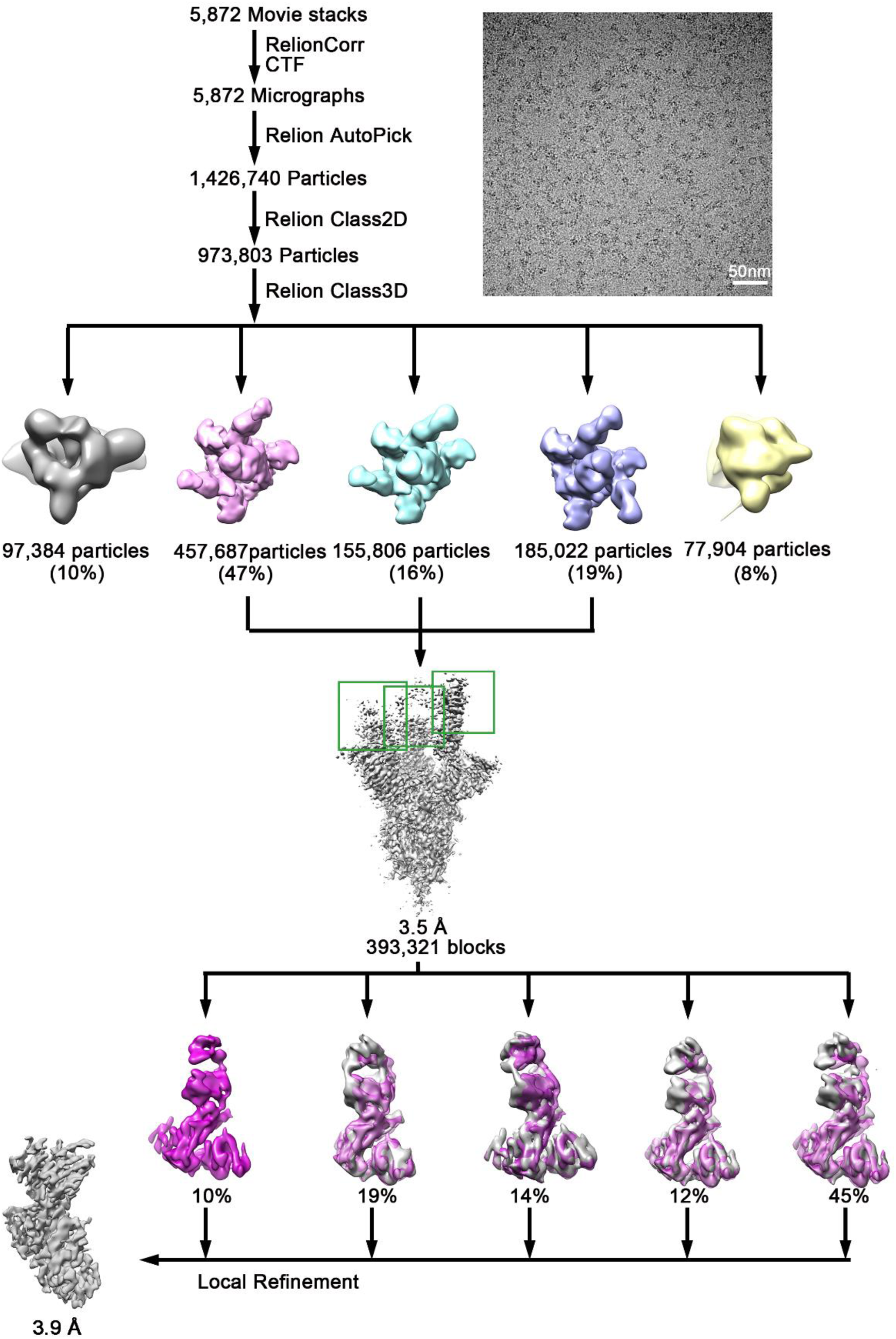
Flowchart of Cryo-EM data processing of SARS-CoV-2 S trimer and HB27 complex. Related to Figure 6.

**Figure S5.**
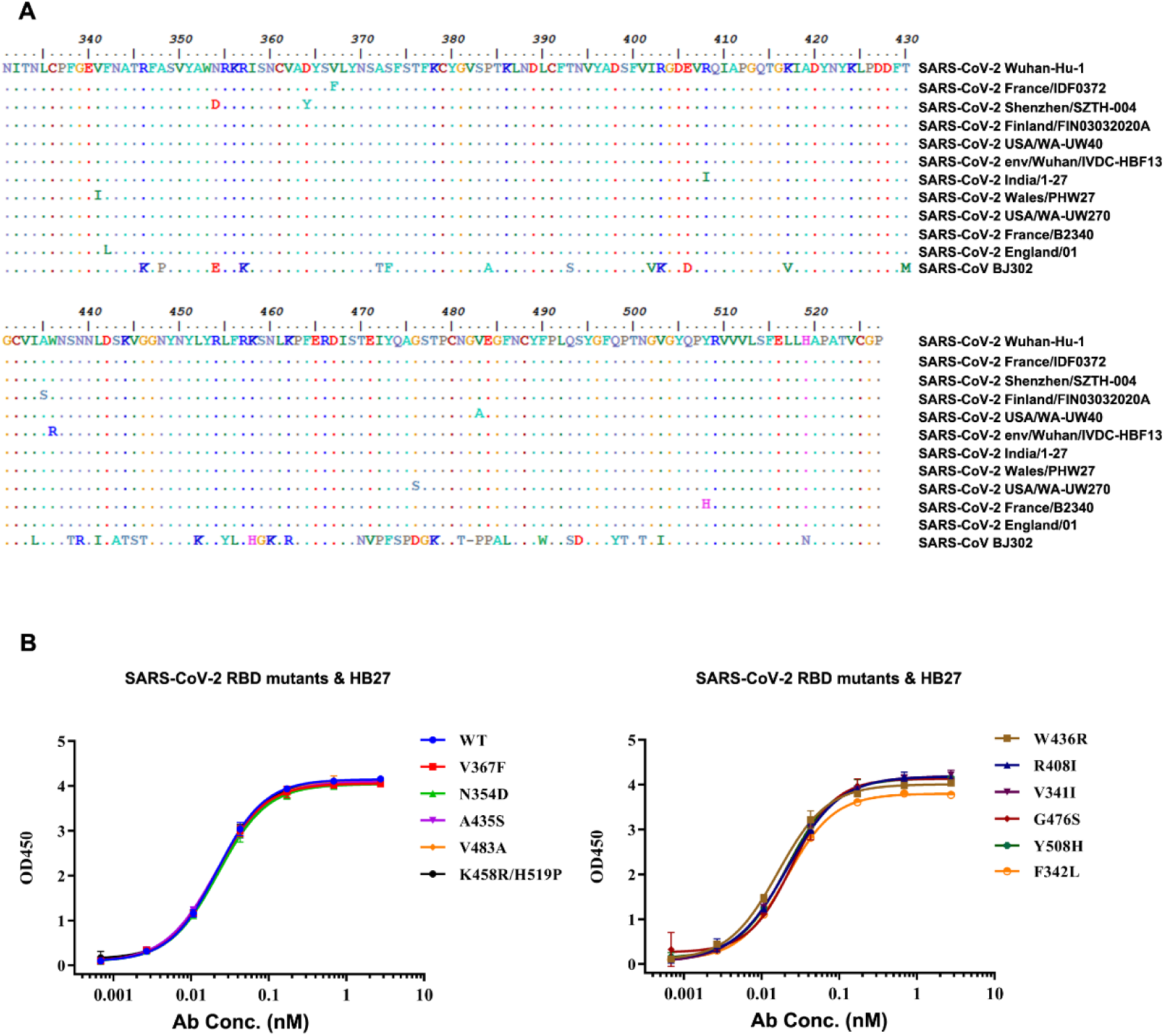
HB27 strongly binds various SARS-CoV-2 RBD mutants. Related to Figure 6. (A) Sequence alignments of the mutated RBDs of circulating SARS-CoV-2 strains used in (A) and SARS-CoV. The genome sequences used in the alignments were downloaded from NCBI and GISAID with accession numbers: NC_045512.2, EPI_ISL_406596, EPI_ISL_406595, EPI_ISL_413602, EPI_ISL_415605, EPI_ISL_408511, EPI_ISL_413522, EPI_ISL_415655, EPI_ISL_418055, EPI_ISL_416507, EPI_ISL_407071 and AY429078.1, respectively. The alignments were analyzed by Clustal W and BioEdit. (B) ELISA binding assays of HB27 with selected SARS-CoV-2 RBD mutants. SARS-CoV-2 RBD proteins with previously reported site mutations were examined for their binding abilities to HB27.

**Figure S6.**
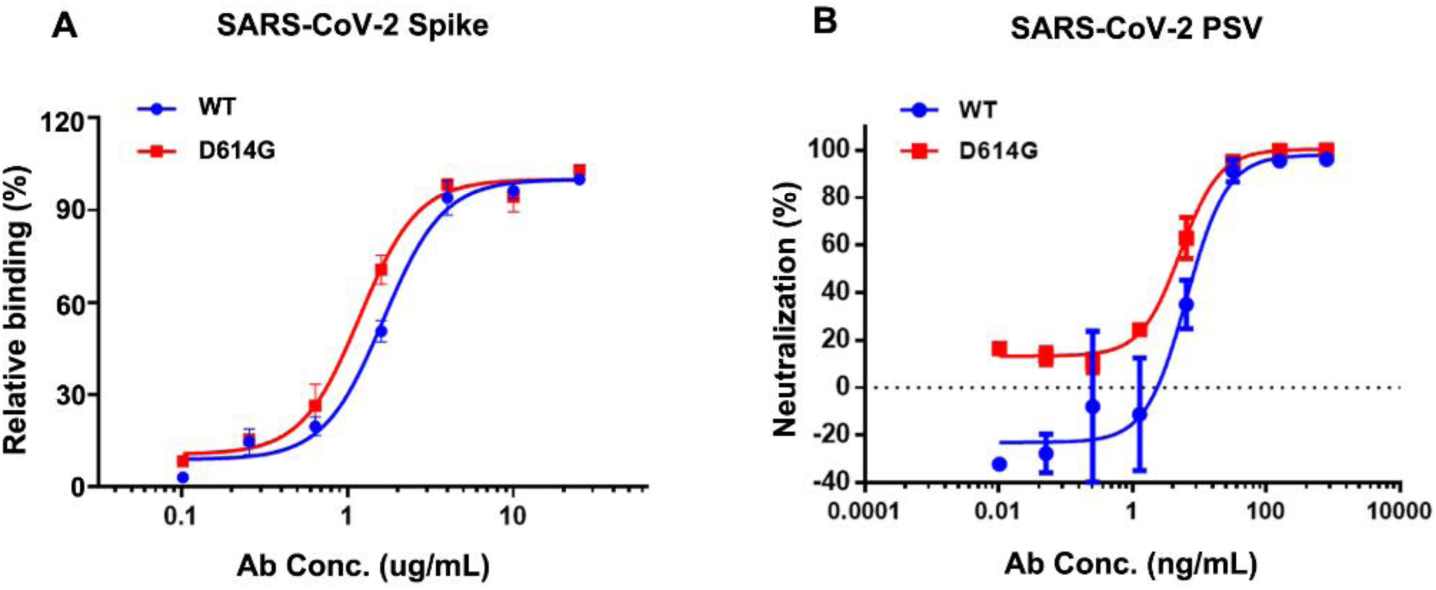
HB27 potently binds and neutralizes SARS-CoV-2 wide type and mutant strain D614G. Related to Figure 7. (A) The spike proteins of WT and D614G were transient expressed in 293T cells which were then examined for binding to HB27 by flow cytometry. (B) Neutralizing activities of HB27 against SARS-CoV-2 WT and D614G pseudoviruses (PSV).

**Figure S7.**
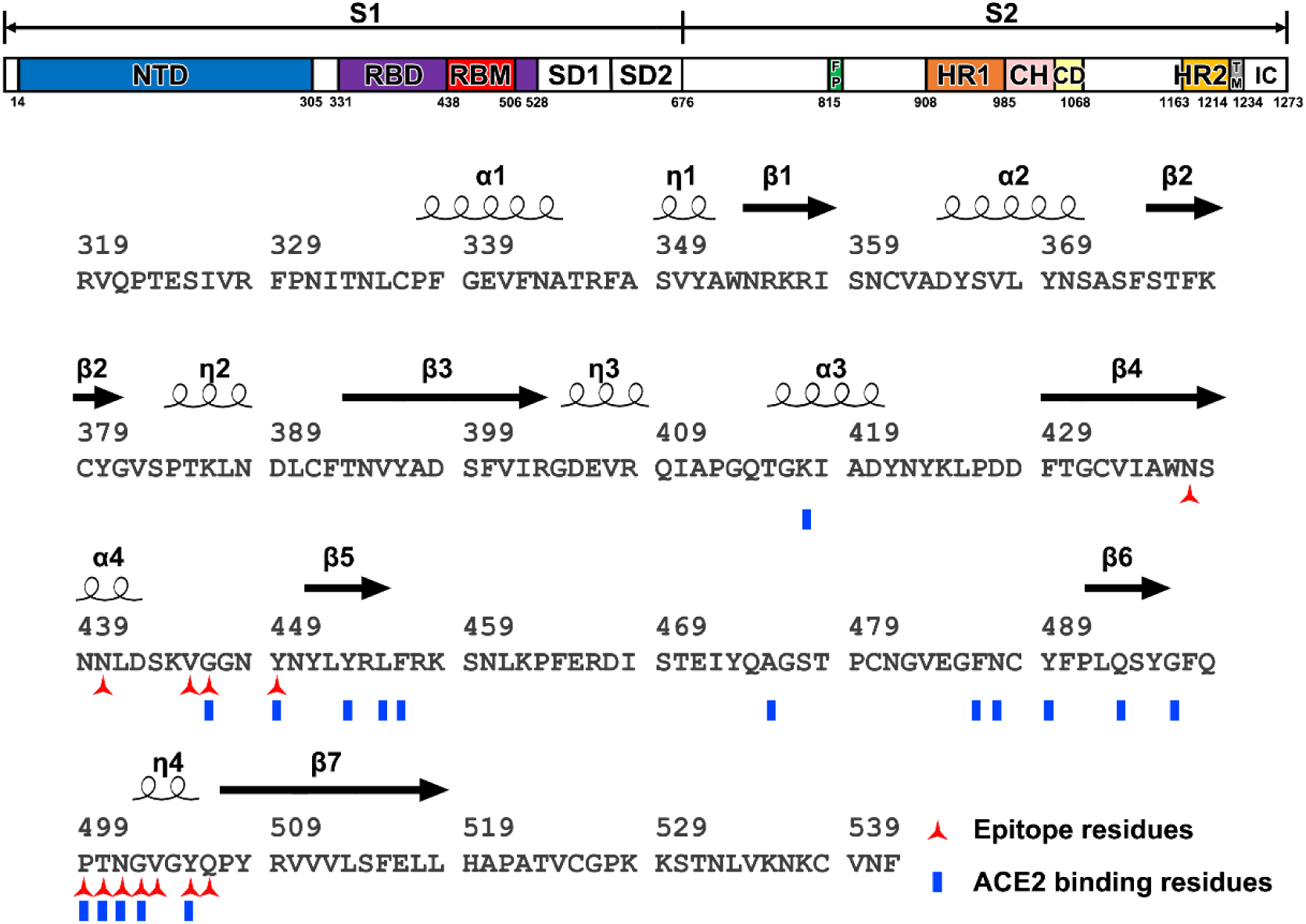
Schematic diagram of SARS-CoV-2 S and the secondary structure of the RBD. Related to Figure 7. (A) Overall topology of SARS-CoV-2 S. NTD: N-terminal domain; RBD: receptor-binding domain; RBM: receptor-binding motif; SD1: subdomain 1; SD2: subdomain 2; FP: fusion peptide; HR1: heptad repeat 1; HR2: heptad repeat 2; TM: transmembrane region; IC: intracellular domain. (B) Protein sequence and the secondary structure of SARS-CoV-2 RBD. The red three-pointed stars and blue rectangles mark the residues in SARS-CoV-2 S RBD that interact with HB27 and ACE2, respectively.

**Table S1.**
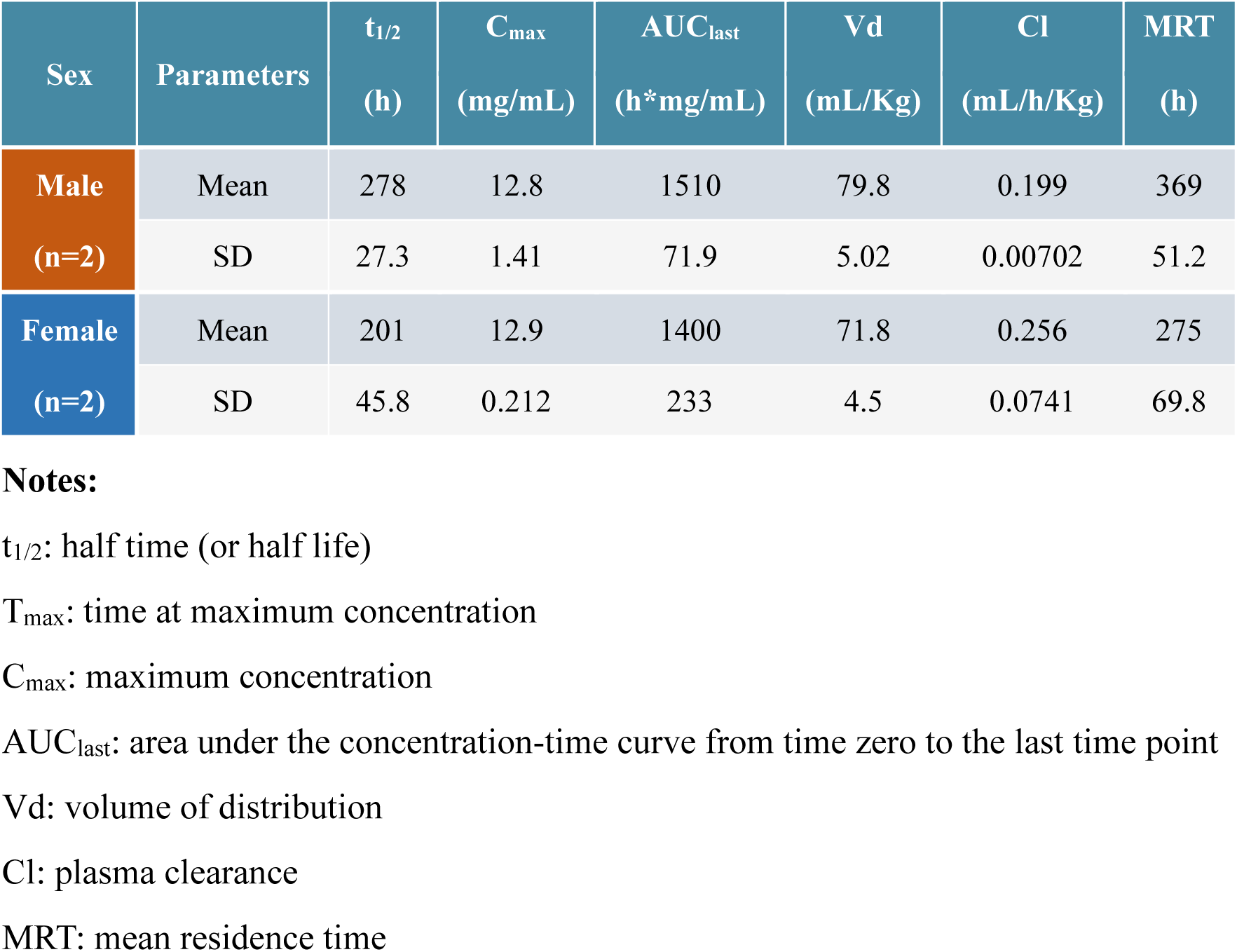
Mean toxicokinetic parameters after intravenous injection of 500 mg/kg HB27 into Rhesus Monkey (0-336 h, n=2, mean ± SD). Related to Figure 3.

| Sex | Parameters | t<sub>1/2</sub><br>(h) | C<sub>max</sub><br>(mg/mL) | AUC<sub>last</sub><br>(h*mg/mL) | Vd<br>(mL/Kg) | Cl<br>(mL/h/Kg) | MRT<br>(h) |
| --- | --- | --- | --- | --- | --- | --- | --- |
| Male<br>(n=2) | Mean | 278 | 12.8 | 1510 | 79.8 | 0.199 | 369 |
|  | SD | 27.3 | 1.41 | 71.9 | 5.02 | 0.00702 | 51.2 |
| Female<br>(n=2) | Mean | 201 | 12.9 | 1400 | 71.8 | 0.256 | 275 |
|  | SD | 45.8 | 0.212 | 233 | 4.5 | 0.0741 | 69.8 |

**Notes:**
- t<sub>1/2</sub>: half time (or half life) - T<sub>max</sub>: time at maximum concentration - C<sub>max</sub>: maximum concentration - AUC<sub>last</sub>: area under the concentration-time curve from time zero to the last time point - Vd: volume of distribution - Cl: plasma clearance - MRT: mean residence time

**Table S2.** Cryo-EM data collection and model refinement statistics. Related to Figures 6.

| **Data collection and reconstruction statistics** |  |  |
| --- | --- | --- |
| Protein | SARS-CoV-2 S-HB27 | Binding interface |
| Voltage (kV) | 300 | 300 |
| Detector | K2 | K2 |
| Pixel size (Å) | 1.04 | 1.04 |
| Electron dose (e<sup>-</sup>/Å<sup>2</sup>) | 60 | 60 |
| Defocus range (μm) | 1.25-2.7 | 1.25-2.7 |
| Final particles | 798,515 | 393,321 |
| Final resolution (Å) | 3.5 | 3.9 |
| **Models refinement and validation statistics** |  |  |
| Ramachandran statistics |  |  |
| Favored (%) | 92.25 | 95.07 |
| Allowed (%) | 6.65 | 3.12 |
| Outliers (%) | 1.09 | 1.81 |
| Rotamer outliers (%) | 0.18 | 0.22 |
| R.m.s.d |  |  |
| Bond lengths (Å) | 0.012 | 0.014 |
| Bond angles (°) | 1.288 | 1.374 |

**Table S3.** Residues of HB27 Fab interacting with the SARS-CoV-2 S trimer at the binding interface (d < 4 Å). Related to Figure 6.

| S-RBD |  | HB27 Fab |  |
| --- | --- | --- | --- |
| Location | Residues | Heavy chain | Light chain |
| **β4** | N437 | G54, G55 |  |
| **α4** | N440 | S52, G55, S56, Y57 |  |
| **α4-β5** | V445 | Y57 |  |
|  | G446 |  | K95 |
|  | Y449 |  | N30, Y31 |
| **β6-η4** | P499 | Y57 |  |
|  | T500 | E50 |  |
|  | N501 | G102 |  |
| **η4** | G502 | N31, Y101 |  |
|  | V503 | N31, S53 |  |
|  | Y505 | Y101, G102 |  |
| **η4-β7** | Q506 | S53 |  |

## References and Notes

Afonine, P.V., Grosse-Kunstleve, R.W., Echols, N., Headd, J.J., Moriarty, N.W., Mustyakimov, M., Terwilliger, T.C., Urzhumtsev, A., Zwart, P.H., and Adams, P.D. (2012). Towards automated crystallographic structure refinement with phenix. refine. Acta Crystallographica Section D: Biological Crystallography 68, 352–367.

Brouwer, P.J.M., Caniels, T.G., van der Straten, K., Snitselaar, J.L., Aldon, Y., Bangaru, S., Torres, J.L., Okba, N.M.A., Claireaux, M., Kerster, G., et al. (2020). Potent neutralizing antibodies from COVID-19 patients define multiple targets of vulnerability. Science.

Brown, A., Long, F., Nicholls, R.A., Toots, J., Emsley, P., and Murshudov, G. (2015). Tools for macromolecular model building and refinement into electron cryo-microscopy reconstructions. Acta Crystallographica Section D-Structural Biology 71, 136–153.

Cao, Y., Su, B., Guo, X., Sun, W., Deng, Y., Bao, L., Zhu, Q., Zhang, X., Zheng, Y., Geng, C., et al. (2020). Potent neutralizing antibodies against SARS-CoV-2 identified by high-throughput single-cell sequencing of convalescent patients’ B cells. Cell.

Chen, V.B., Arendall, W.B., Headd, J.J., Keedy, D.A., Immormino, R.M., Kapral, G.J., Murray, L.W., Richardson, J.S., and Richardson, D.C. (2010). MolProbity: all-atom structure validation for macromolecular crystallography. Acta Crystallographica Section D: Biological Crystallography 66, 12–21.

Corti, D., Zhao, J., Pedotti, M., Simonelli, L., Agnihothram, S., Fett, C., Fernandez-Rodriguez, B., Foglierini, M., Agatic, G., Vanzetta, F., et al. (2015). Prophylactic and postexposure efficacy of a potent human monoclonal antibody against MERS coronavirus. Proceedings of the National Academy of Sciences of the United States of America 112, 10473–10478.

Du, L., He, Y., Zhou, Y., Liu, S., Zheng, B.J., and Jiang, S. (2009). The spike protein of SARS-CoV—a target for vaccine and therapeutic development. Nature reviews Microbiology 7, 226–236.

Gallagher, T.M., and Buchmeier, M.J. (2001). Coronavirus spike proteins in viral entry and pathogenesis. Virology 279, 371–374.

Gao, Q., Bao, L., Mao, H., Wang, L., Xu, K., Yang, M., Li, Y., Zhu, L., Wang, N., Lv, Z., et al. (2020). Rapid development of an inactivated vaccine candidate for SARS-CoV-2. Science.

Gu, H., Chen, Q., Yang, G., He, L., Fan, H., Deng, Y.-Q., Wang, Y., Teng, Y., Zhao, Z., Cui, Y., et al. (2020). Rapid adaptation of SARS-CoV-2 in BALB/c mice: Novel mouse model for vaccine efficacy. bioRxiv, 2020. 2005.2002.073411.

Gui, M., Song, W., Zhou, H., Xu, J., Chen, S., Xiang, Y., and Wang, X. (2017). Cryo-electron microscopy structures of the SARS-CoV spike glycoprotein reveal a prerequisite conformational state for receptor binding. Cell research 27, 119–129.

Hansen, J., Baum, A., Pascal, K.E., Russo, V., Giordano, S., Wloga, E., Fulton, B.O., Yan, Y., Koon, K., Patel, K., et al. (2020). Studies in humanized mice and convalescent humans yield a SARS-CoV-2 antibody cocktail. Science.

Hoffmann, M., Kleine-Weber, H., Schroeder, S., Kruger, N., Herrler, T., Erichsen, S., Schiergens, T.S., Herrler, G., Wu, N.H., Nitsche, A., et al. (2020). SARS-CoV-2 Cell Entry Depends on ACE2 and TMPRSS2 and Is Blocked by a Clinically Proven Protease Inhibitor. Cell 181, 271–280 e278.

Kirchdoerfer, R.N., Cottrell, C.A., Wang, N., Pallesen, J., Yassine, H.M., Turner, H.L., Corbett, K.S., Graham, B.S., McLellan, J.S., and Ward, A.B. (2016). Pre-fusion structure of a human coronavirus spike protein. Nature 531, 118–121.

Korber, B., Fischer, W., Gnanakaran, S.G., Yoon, H., Theiler, J., Abfalterer, W., Foley, B., Giorgi, E.E., Bhattacharya, T., and Parker, M.D. (2020). Spike mutation pipeline reveals the emergence of a more transmissible form of SARS-CoV-2. bioRxiv : the preprint server for biology.

Kucukelbir, A., Sigworth, F.J., and Tagare, H.D. (2014). Quantifying the local resolution of cryo-EM density maps. Nature methods 11, 63–65.

Li, F. (2016). Structure, Function, and Evolution of Coronavirus Spike Proteins. Annual review of virology 3, 237–261.

Lu, R., Zhao, X., Li, J., Niu, P., Yang, B., Wu, H., Wang, W., Song, H., Huang, B., Zhu, N., et al. (2020). Genomic characterisation and epidemiology of 2019 novel coronavirus: implications for virus origins and receptor binding. Lancet 395, 565–574.

Nie, J., Li, Q., Wu, J., Zhao, C., Hao, H., Liu, H., Zhang, L., Nie, L., Qin, H., Wang, M., et al. (2020). Establishment and validation of a pseudovirus neutralization assay for SARS-CoV-2. Emerging microbes & infections 9, 680–686.

Ou, X., Liu, Y., Lei, X., Li, P., Mi, D., Ren, L., Guo, L., Guo, R., Chen, T., Hu, J., et al. (2020). Characterization of spike glycoprotein of SARS-CoV-2 on virus entry and its immune cross-reactivity with SARS-CoV. Nature communications 11, 1620.

Pallesen, J., Wang, N., Corbett, K.S., Wrapp, D., Kirchdoerfer, R.N., Turner, H.L., Cottrell, C.A., Becker, M.M., Wang, L., Shi, W., et al. (2017). Immunogenicity and structures of a rationally designed prefusion MERS-CoV spike antigen. Proceedings of the National Academy of Sciences of the United States of America 114, E7348–E7357.

Pettersen, E.F., Goddard, T.D., Huang, C.C., Couch, G.S., Greenblatt, D.M., Meng, E.C., and Ferrin, T.E. (2004). UCSF Chimera—a visualization system for exploratory research and analysis. Journal of computational chemistry 25, 1605–1612.

Pinto, D., Park, Y.J., Beltramello, M., Walls, A.C., Tortorici, M.A., Bianchi, S., Jaconi, S., Culap, K., Zatta, F., De Marco, A., et al. (2020). Cross-neutralization of SARS-CoV-2 by a human monoclonal SARS-CoV antibody. Nature.

Qiu, X., Lei, Y., Yang, P., Gao, Q., Wang, N., Cao, L., Yuan, S., Huang, X., Deng, Y., Ma, W., et al. (2018). Structural basis for neutralization of Japanese encephalitis virus by two potent therapeutic antibodies. Nature microbiology 3, 287–294.

Scheres, S.H. (2016). Processing of Structurally Heterogeneous Cryo-EM Data in RELION. Methods in enzymology 579, 125–157.

Scheres, S.H., and Chen, S. (2012). Prevention of overfitting in cryo-EM structure determination. Nature methods 9, 853–854.

Shang, J., Wan, Y., Luo, C., Ye, G., Geng, Q., Auerbach, A., and Li, F. (2020). Cell entry mechanisms of SARS-CoV-2. Proceedings of the National Academy of Sciences of the United States of America 117, 11727–11734.

Shi, R., Shan, C., Duan, X., Chen, Z., Liu, P., Song, J., Song, T., Bi, X., Han, C., Wu, L., et al. (2020). A human neutralizing antibody targets the receptor binding site of SARS-CoV-2. Nature.

Sun, S.H., Chen, Q., Gu, H.J., Yang, G., Wang, Y.X., Huang, X.Y., Liu, S.S., Zhang, N.N., Li, X.F., Xiong, R., et al. (2020). A Mouse Model of SARS-CoV-2 Infection and Pathogenesis. Cell host & microbe.

Walls, A.C., Park, Y.J., Tortorici, M.A., Wall, A., McGuire, A.T., and Veesler, D. (2020). Structure, Function, and Antigenicity of the SARS-CoV-2 Spike Glycoprotein. Cell 181, 281–292 e286.

Walls, A.C., Tortorici, M.A., Snijder, J., Xiong, X., Bosch, B.J., Rey, F.A., and Veesler, D. (2017). Tectonic conformational changes of a coronavirus spike glycoprotein promote membrane fusion. Proceedings of the National Academy of Sciences of the United States of America 114, 11157–11162.

Walls, A.C., Xiong, X., Park, Y.J., Tortorici, M.A., Snijder, J., Quispe, J., Cameroni, E., Gopal, R., Dai, M., Lanzavecchia, A., et al. (2019). Unexpected Receptor Functional Mimicry Elucidates Activation of Coronavirus Fusion. Cell 176, 1026–1039 e1015.

Wang, N., Chen, W., Zhu, L., Zhu, D., Feng, R., Wang, J., Zhu, B., Zhang, X., Chen, X., Liu, X., et al. (2020). Structures of the portal vertex reveal essential protein-protein interactions for Herpesvirus assembly and maturation. Protein & cell 11, 366–373.

Wang, N., Zhao, D., Wang, J., Zhang, Y., Wang, M., Gao, Y., Li, F., Wang, J., Bu, Z., Rao, Z., et al. (2019). Architecture of African swine fever virus and implications for viral assembly. Science 366, 640–644.

Wang, X., Zhu, L., Dang, M., Hu, Z., Gao, Q., Yuan, S., Sun, Y., Zhang, B., Ren, J., Kotecha, A., et al. (2017). Potent neutralization of hepatitis A virus reveals a receptor mimic mechanism and the receptor recognition site. Proceedings of the National Academy of Sciences of the United States of America 114, 770–775.

Wec, A.Z., Wrapp, D., Herbert, A.S., Maurer, D.P., Haslwanter, D., Sakharkar, M., Jangra, R.K., Dieterle, M.E., Lilov, A., Huang, D., et al. (2020). Broad neutralization of SARS-related viruses by human monoclonal antibodies. Science.

Wrapp, D., Wang, N., Corbett, K.S., Goldsmith, J.A., Hsieh, C.L., Abiona, O., Graham, B.S., and McLellan, J.S. (2020). Cryo-EM structure of the 2019-nCoV spike in the prefusion conformation. Science 367, 1260–1263.

Wu, Y., Wang, F., Shen, C., Peng, W., Li, D., Zhao, C., Li, Z., Li, S., Bi, Y., Yang, Y., et al. (2020). A noncompeting pair of human neutralizing antibodies block COVID-19 virus binding to its receptor ACE2. Science.

Xia, S., Liu, M., Wang, C., Xu, W., Lan, Q., Feng, S., Qi, F., Bao, L., Du, L., Liu, S., et al. (2020). Inhibition of SARS-CoV-2 (previously 2019-nCoV) infection by a highly potent pan-coronavirus fusion inhibitor targeting its spike protein that harbors a high capacity to mediate membrane fusion. Cell research 30, 343–355.

Yang, Y., Yang, P., Wang, N., Chen, Z., Su, D., Zhou, Z.H., Rao, Z., and Wang, X. (2020). Architecture of the herpesvirus genome-packaging complex and implications for DNA translocation. Protein & cell 11, 339–351.

Yuan, M., Wu, N.C., Zhu, X., Lee, C.D., So, R.T.Y., Lv, H., Mok, C.K.P., and Wilson, I.A. (2020). A highly conserved cryptic epitope in the receptor binding domains of SARS-CoV-2 and SARS-CoV. Science 368, 630–633.

Zhang, K. (2016). Gctf: Real-time CTF determination and correction. Journal of structural biology 193, 1–12.

Zhe Lv, Y.-Q.D., Qing Ye, Lei Cao, Chun-Yun Sun, Changfa Fan, Weijin Huang, Shihui Sun, Yao Sun, Ling Zhu, Qi Chen, Nan Wang, Jianhui Nie, Zhen Cui, Dandan Zhu, Neil Shaw, Xiao-Feng Li, Qianqian Li, Liangzhi Xie, Youchun Wang, Zihe Rao, Cheng-Feng Qin, Xiangxi Wang (2020). Structural basis for neutralization of SARS-CoV-2 and SARS-CoV by a potent therapeutic antibody. Science.

Zhou, P., Yang, X.L., Wang, X.G., Hu, B., Zhang, L., Zhang, W., Si, H.R., Zhu, Y., Li, B., Huang, C.L., et al. (2020). A pneumonia outbreak associated with a new coronavirus of probable bat origin. Nature 579, 270–273.

Zost, S.J., Gilchuk, P., Case, J.B., Binshtein, E., Chen, R.E., Nkolola, J.P., Schafer, A., Reidy, J.X., Trivette, A., Nargi, R.S., et al. (2020). Potently neutralizing and protective human antibodies against SARS-CoV-2. Nature 584, 443–449.

